# Synergistic cross-kingdom host cell damage between *Candida albicans* and *Enterococcus faecalis*

**DOI:** 10.1101/2024.09.11.612452

**Authors:** Mario Kapitan, Maria Joanna Niemiec, Nicolas Millet, Philipp Brandt, Md Estiak Khan Chowdhury, Anna Czapka, Ketema Abdissa, Franziska Hoffmann, Anna Lange, Mark Veleba, Sandor Nietzsche, Alexander S. Mosig, Bettina Löffler, Mike Marquet, Oliwia Makarewicz, Kimberly A. Kline, Slavena Vylkova, Marc Swidergall, Ilse D. Jacobsen

## Abstract

The fungus *Candida albicans* and the Gram-positive bacterium *Enterococcus faecalis* share mucosal niches in the human body. As opportunistic pathogens, both are found to expand population size during dysbiosis, and can cause severe systemic infections in susceptible individuals. Here, we show that the presence of *C. albicans* results in increased host cell damage by *E. faecalis*. Furthermore, *E. faecalis* aggravates oropharyngeal candidiasis in mice. Increased damage is mediated by enterococcal cytolysin, and involves both physical interaction and altered glucose availability. Physical interaction promotes accumulation of bacteria on host cells, facilitating contact of cytolysin with host cells. Glucose depletion by the metabolic activity of the fungus sensitized host cells to cytolysin. This work illustrates how a complex interplay between fungi and bacteria can result in detrimental consequences for the host.

## Introduction

*Candida albicans* is a fungal pathobiont colonizing different mucosal surfaces of the human body. Colonization is asymptomatic in healthy individuals, but translocation across mucosal barriers in hospitalized and/or immunocompromised patients can lead to severe systemic infection (McCarty and Pappas, 2016; Pappas et al., 2003). Besides immunosuppression, disturbance of the mucosal microbiome by antibiotic treatment is a major risk factor for disseminated candidiasis (Pappas et al., 2003), suggesting that fungal-bacterial interactions significantly influence the behaviour of *C. albicans* in the host. While most research has focused on antagonistic effects of bacteria on *C. albicans* virulence or colonization, synergism has been observed for a few bacterial species, including *Enterococcus faecalis* (Bertolini et al., 2021b; Klaerner et al., 1997; Mason et al., 2012a; Mason et al., 2012b; Todd et al., 2019b).

Enterococci are lactic acid producing Gram positive bacteria ubiquitous in nature and colonizers of plants and animals, including humans (Mundt, 1963a; Mundt, 1963b; Noble, 1978). Like *C. albicans*, they are opportunistic pathogens that can cause various infections ranging from urinary tract infections to bacteraemia and endocarditis (Graninger and Ragette, 1992; Hidron et al., 2008). They share various mucosal host niches with *C. albicans*, like the oral cavity (Smyth et al., 1987), urogenital (Kline and Lewis, 2016) and the intestinal tract (Noble, 1978). In the gut and the oral cavity, the presence of *C. albicans* increases the relative abundance of *Enterococcus* spp. after perturbation of the mucosal microbiota by antibiotics (Erb Downward et al., 2013; Mason et al., 2012a; Mason et al., 2012b) or chemotherapy (Bertolini et al., 2019). Furthermore, *Candida* and enterococci are commonly co-isolated from clinical samples, including blood cultures (Hermann et al., 1999; Klotz et al., 2007; Sutter et al., 2008).

While these observations suggest synergistic interactions between *C. albicans* and enterococci, with possible clinical consequences, *E. faecalis* has also been shown to reduce *C. albicans* virulence by secretion of the bacteriocin EntV. Therefore, it was postulated that *E. faecalis* promotes a commensal lifestyle of *C. albicans* by inhibiting fungal filamentation, which in turn would benefit *E. faecalis* by synergistic interactions (Garsin and Lorenz, 2013). Contrary to this hypothesis, however, interaction of *E. faecalis* and *C. albicans* on an oral organoid model led to increased tissue erosion and microbial invasion (Krishnamoorthy et al., 2020). In this model, virulence-associated *E. faecalis* genes were downregulated, whereas fungal virulence-associated genes were upregulated.

One limitation of the previous studies on *C. albicans* – *E. faecalis* interactions is that only few strains were used. Enterococci are known for high genome plasticity, with a small core genome and a large accessory genome contributing to up to 25% of the total genome size (Cattoir, 2022). Horizontal gene transfer is common in enterococci, involves different types of mobile genetic elements, and has mainly been studied in the context of antimicrobial resistance (Johnson et al., 2021). In addition, the *E. faecalis* pathogenicity island can be horizontally transferred (Johnson et al., 2021) and contains variations between strains that modulate virulence (Shankar et al., 2002). To address the possible impact of strain-to-strain variation of *E. faecalis* on interaction with *C. albicans*, we screened a larger number of strains in an *in vitro* intestinal model. We discovered that the type of bacterial-fungal interaction depends on strain-specific properties of the bacterial partner. Specifically, synergistic host cell damage depended on the *E. faecalis* cytolysin. We confirmed the *in vivo* relevance of these findings by using a murine model of oral candidiasis. In this model, cytolysin-producing *E. faecalis* aggravated invasion and damage, while a cytolysin-negative EntV-producing strain reduced invasion. The presence of *C. albicans* increased bacterial virulence partly by increasing *E. faecalis* accumulation and aggregation on host cells. In addition, fungal-mediated nutrient depletion sensitized host cells to pore-forming toxins *in vitro*. Thus, multiple mechanisms contribute to synergistic damage caused *C. albicans*-*E. faecalis* coinfection.

## Results

### Host cell damage after coinfection of an *in vitro* intestinal model with *Candida albicans* and *Enterococcus faecalis* is bacterial strain-dependent

Both antagonistic and synergistic interactions between *C. albicans* and *E. faecalis* have been reported (Bertolini et al., 2019; Cruz et al., 2013; Graham et al., 2017; Hermann et al., 1999; Klotz et al., 2007; Mason et al., 2012b). To determine the type of interaction on enterocytes, we used an *in vitro* cell culture model to quantify host cell damage following either single infection with *C. albicans* or enterococci, and damage after coinfection. The *E. faecalis* type strain, the commonly used lab strains ATCC 29212, OG1RF, and V583, as well as 16 clinical isolates obtained from blood, stool, or urine were analysed to address possible variations between bacterial strains. While for most of the tested *E. faecalis* isolates the coinfection damage was comparable to or slightly lower than the sum of damage caused by fungus and bacteria alone, five *E. faecalis* strains displayed significantly higher coinfection damage under ambient oxygen (21%), reduced oxygen (10%), and low but physiological gut oxygen tension (1% O_2_; Fig. 1A-C). The synergistic effect was especially pronounced at medium and low oxygen. By quantification of bacterial and fungal burden, we could exclude that enhanced host cell damage was due to increased proliferation of microbes during coinfection (Fig. S1).

**Figure 1:**
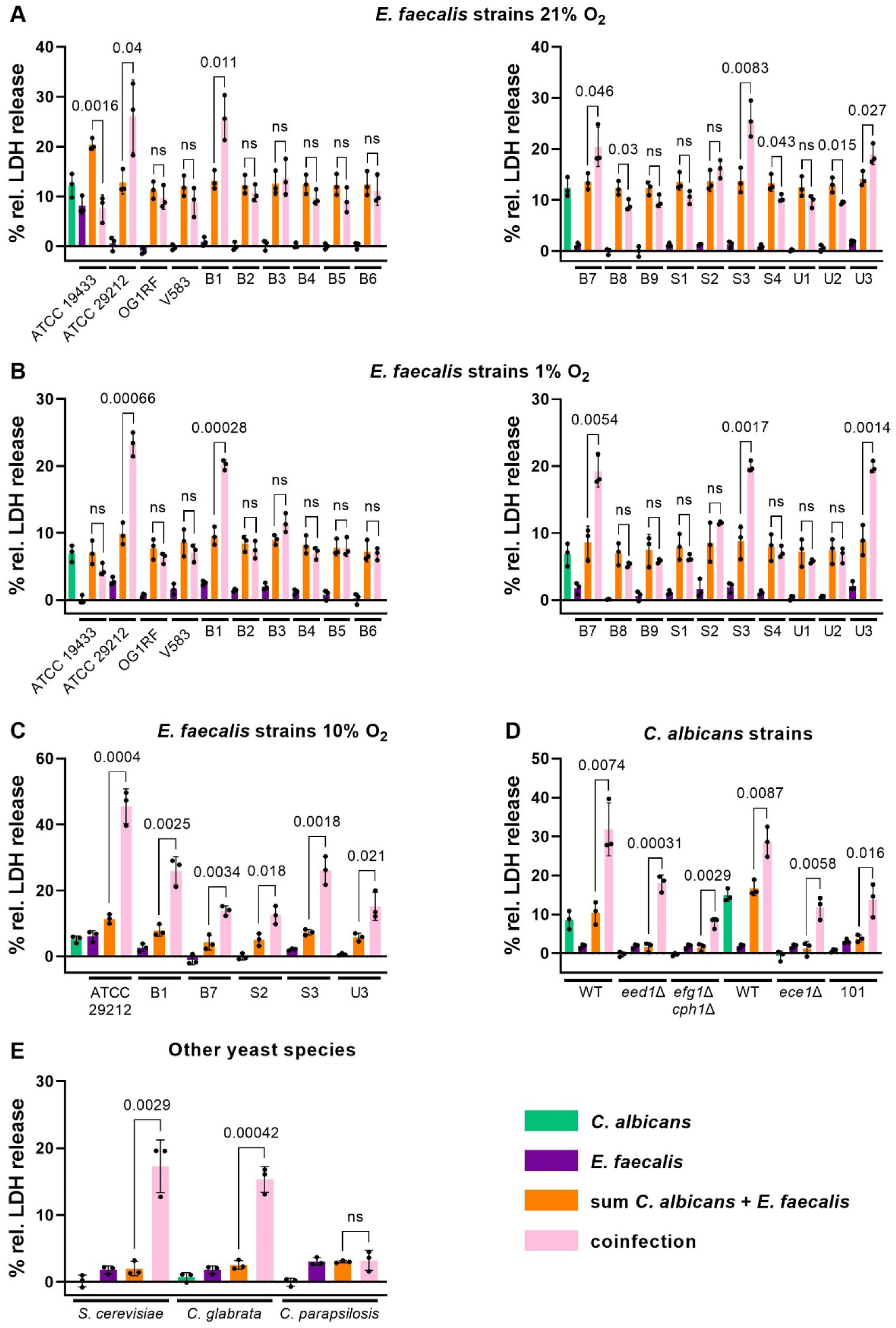
Strain-dependent synergy during *Candida*-*Enterococcus* coinfection of enterocytes results in increased host cell damage. Enterocytes were infected with 10^6^ cells/ml *C. albicans* for 6 h followed by addition of 10^5^ cells/ml *E. faecalis* and further incubation for 24 h. Damage was measured by release of host cell lactate dehydrogenase (LDH) into the supernatant, normalised to non-infected controls, and is displayed as percent damage relative to Triton X-100-lysis. Green: monoinfection with *C. albicans*; dark purple: monoinfection with *E. faecalis*; orange: sum of single infections; light pink: coinfection. Bars represent mean and SD, n = 3. The sum of LDH release caused by bacterial and fungal monoinfection was compared to the LDH release following coinfection using two-tailed unpaired Student’s t-tests. Significant values are indicated by p-values. ns: not significant (p > 0.05). (**A**) Damage after infection with *C. albicans* SC5314 and different *E. faecalis* strains at 21 % O_2_, 5 % CO_2_. (**B**) Damage at 1 % O_2,_ 5 % CO_2_. **C:** Damage at 10 % O_2,_ 5 % CO_2_. (**D, E**) Damage after infection with *E. faecalis* B1 and different *C. albicans* strains (**D**) or other yeast species WT: Parental strain of the respective deletion mutants (SC5314 for *eed1*Δ and *efg1*Δ*cph1*Δ; BWP17 for *ece1*Δ).

To determine if the overall higher microbial burden during coinfection led to increased host cell damage, we tested if the number of bacterial and fungal cells used for infection affects LDH release from host cells. Increasing the bacterial inoculum over a range of 4 log_10_ led to only a twofold increase in monoinfection damage, and coinfection damage was similar within a range of 10^4^ to 10^6^ bacteria per ml (Fig. S2A). Similarly, *C. albicans*-induced damage was relatively stable in monoinfections; however, coinfection damage was concentration-dependent (Fig. S2B). One possible explanation for the lack of a clear dose-dependent increase of enterocyte damage in mono-infections is that bacterial growth in the system is limited, as suggested by the lack of further increase of bacterial CFU from 24 to 30h shown in Fig. S1. For *C. albicans*, high numbers of fungal filaments might multiple layers of fungal cells without increased numbers of fungal filaments accessing and damaging enterocytes. This indicated that synergistic host cell damage could not be explained by the higher microbial burden, but suggested that fungal activities were required. Consistent with this, heat-killed *C. albicans* did not induce synergistic host cell damage during coinfection (Fig. S2C). Likewise, bacterial viability was required for inducing increased damage (Fig. S2D), indicating that both microbes contribute to the synergistic virulence phenotype. Furthermore, synergistic damage developed within the last 6 h of the co-incubation period, in which bacterial CFU exceeded fungal CFU > 100fold (Fig. S1) and damage caused by monoinfections remained stable (Fig. S2E).

### *C. albicans* virulence factors contribute to damage during coinfection but are not required for synergism

In the experiments presented above, enterocytes were infected with *C. albicans* first, and bacteria were added after 6 h to prevent a possible negative impact of fast bacterial replication on fungal growth or germ tube induction at early stages of infection. We next tested if this preincubation was required for synergistic damage, and observed that synergism became evident and more pronounced with prolonged preincubation of host cells with *C. albicans* (Fig. S2F).

*C. albicans* rapidly forms germ tubes and hyphae in cell culture conditions, and thereby could contribute to synergistic damage. To test this, we used *C. albicans* mutants with defects in filamentation (*eed1*Δ/*eed1*Δ and *efg1*Δ*cph1*Δ/*efg1*Δ*cph1*Δ). As expected, these mutants were strongly reduced in their ability to damage host cells during monoinfection (Fig. 1D). These strains were still able to induce synergistic damage during coinfection. Synergistic damage was also observed with a mutant lacking *ECE1* encoding the fungal peptide toxin candidalysin, and the commensal *C. albicans* strain 101 (Fig. 1D). Overall coinfection damage with was lower for the *C. albicans* mutants and 101 compared to the virulent wildtype strains; this is likely due to the lower or absent fungal-mediated damage in coinfection with these strains. To determine if the synergistic effect on coinfection damage is limited to *C. albicans*, we tested additional fungal species: *Saccharomyces cerevisiae* and *Candida glabrata*, but not *Candida parapsilosis*, induced increased coinfection damage (Fig. 1E) despite their lack of cytotoxicity in mono-infections. Taken together, these findings suggested that the increased host cell damage during coinfection was mainly caused by bacterial activities.

### Enterococcal cytolysin is necessary for enhanced coinfection damage of enterocytes

Several *E. faecalis* factors contributing to virulence and host cell damage have been described (Fisher and Phillips, 2009). We hypothesized that synergism with fungal infection is mediated by a distinct virulence factor, or set of factors, found in all synergistic strains, but absent in non-synergistic isolates. Therefore, we screened the genomes of the *E. faecalis* isolates used for the presence of virulence-associated genes. Synergistic strains belong to different capsule operon types, but share several virulence-associated genes, e.g. encoding aggregation substance, certain pili, adhesins, and gelatinase (Table S1). However, this set of genes was also present in non-synergistic strains, including OG1RF (Table S2). A notable exception was the cytolysin operon (Table S3). Cytolysin is a pore-forming toxin consisting of two peptides and causing haemolysis on horse blood agar plates (HBA). It is an important virulence factor of *E. faecalis* causing host cell damage, and is associated with increased virulence in infection models and patient mortality (Garsin et al., 2001; Huycke et al., 1991; Ike et al., 1984; Jett et al., 1995; Jett et al., 1992). All synergistic strains displayed β-haemolysis on HBA, and carried the complete cytolysin operon (*cyl*; Table S3). The two strains that appear to contain the full *cyl* operon but displayed no synergism in co-infections, strain B3 and B8, showed no β-haemolysis on HBA at high oxygen, and only B3 displayed weak haemolysis at low oxygen (Table S3), indicating possible defects in cytolysin regulation or expression.

The *cyl* operon is often located on self-transmissible plasmids (Jett et al., 1992), and sequencing results indicated the presence of a 64 kb plasmid containing the cyl operon in the synergistic strain B1 (Fig. S3). To explore the possible role of cytolysin for synergistic damage, we therefore performed plasmid curing of strain B1 and selected haemolytic and non-haemolytic derivative strains for coinfection experiments. The loss of haemolytic activity was associated with the loss of increased coinfection damage (Fig. 2A). Conversely, transmission of haemolytic activity from B1 to the non-synergistic plasmid-free strain OG1RF by conjugation resulted in increased host cell damage during coinfection (Fig. 2B; Fig. S2G, H). This suggested the presence of a conjugative plasmid in B1, and a link between haemolysis and synergistic damage. However, we cannot exclude that some of the haemolytic colonies obtained in the conjugation experiment were spontaneous rifampicin-resistant mutants of the donor strain. Furthermore, other factors encoded on the putative plasmid could have contributed to the phenotype. We therefore analysed isogenic mutants of *E. faecalis* OG1X (Ehrenfeld and Clewell, 1987; Ike and Clewell, 1984; Ike et al., 1990; Ike et al., 1983) harbouring plasmids encoding the cytolysin operon and/or the gene encoding for the adhesin aggregation substance (AS) (Kreft et al., 1992), which increases adhesion and internalization of *E. faecalis* by colonic enterocytes (Sartingen et al., 2000). Increased damage of host cells during coinfection was consistently observed for cytolysin positive strains (Fig. 2C), while the presence of AS had no effect on damage. Thus, enterococcal cytolysin is required for the enhanced damage during coinfection, and the presence or absence of the *cyl* operon explains the differences in coinfection damage observed for the different *E. faecalis* strains.

**Figure 2:**
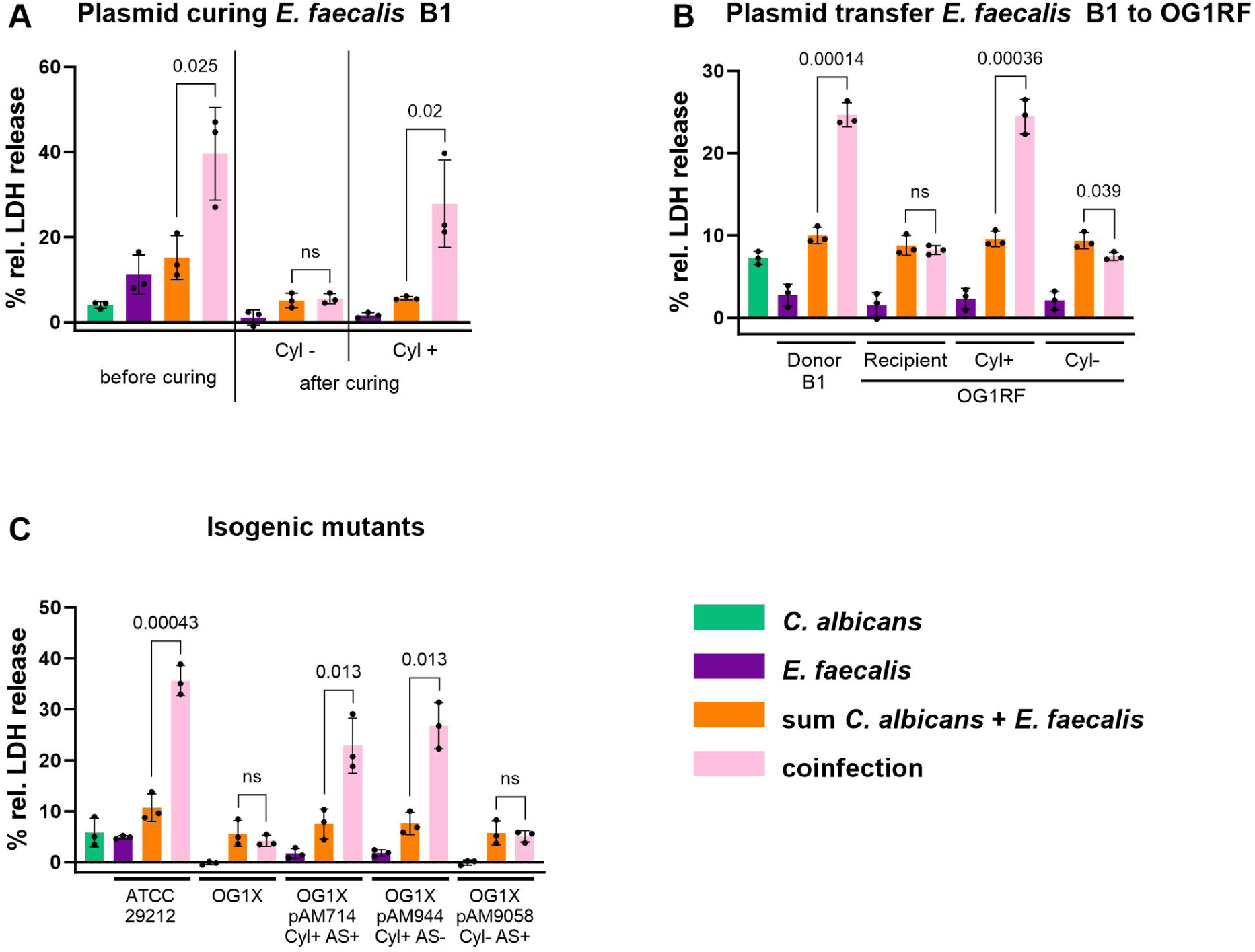
Synergistic coinfection damage is mediated by the *E. faecalis* cytolysin. Enterocytes were infected with 10^6^ cells/ml *C. albicans* SC5314 for 6 h at 1 % O_2_ followed by addition of 10^5^ cells/ml *E. faecalis* and further incubation for 24 h. Damage was measured by release of host cell lactate dehydrogenase (LDH) into the supernatant, normalised to non-infected controls, and is displayed as percent damage relative to Triton X-100-lysis Green: monoinfection with *C. albicans*; dark purple: monoinfection with *E. faecalis*; orange: sum of single infections; light pink: coinfection. Bars represent mean and SD, n = 3. The sum of LDH release caused by bacterial and fungal monoinfection was compared to the LDH release following coinfection using two-tailed unpaired Student’s t-tests. Significant values are indicated by p-values. ns: not significant (p > 0.05). (**A**) Loss of haemolytic activity after plasmid curing of *E. faecalis* B1 coincides with loss of synergistic coinfection damage. Cyl-: non-haemolytic *E. faecalis* B1 obtained by plasmid curing; Cyl+: *E. faecalis* B1 isolated obtained from the same plasmid curing approach which retained haemolytic activity. (**B**) Transfer of haemolytic activity from *E. faecalis* B1 (donor) to *E. faecalis* OG1RF (recipient) by conjugation results in synergistic coinfection damage. Cyl+: haemolytic OG1RF derivative; Cyl-: non-haemolytic OG1RF derivative obtained from the same conjugation experiment. (**C**) Coinfection with OG1X (plasmid free) and OG1X derivatives harbouring the following plasmids: pAM714: cytolysin-positive/agglutination substance-positive; pAM944: cytolysin-positive/agglutination substance-negative; pAM9058: cytolysin-negative/agglutination substance-positive) and ATCC 29212 (haemolytic) as control. Synergistic coinfection coincides with a functional cytolysin operon.

### The effect of *E. faecalis* on the severity of murine oropharyngeal candidiasis depends on the bacterial cytolysin

*C. albicans* and *E. faecalis* do not only share a niche in the gastrointestinal tract but also in the oral cavity. We therefore quantified host cell damage during mono- and coinfection of oral buccal cells (TR146). For these experiments, we used an MOI of 1 to achieve a similar level of damage with *C. albicans* alone as observed with enterocytes. Similar to intestinal cells, coinfection damage was increased with cytolysin positive strains of *E. faecalis* (Fig. 3A). To determine if these strain-dependent effects can be translated to *in vivo* infections, we used a murine oropharyngeal candidiasis (OPC) model in which the drinking water was supplemented with different *E. faecalis* strains after *C. albicans* infection. Since we observed apoptotic cell death following infection *in vitro* both for intestinal and epithelial cells (Fig. S4), apoptosis was included as read out for *in vivo* invasion in addition to invasion depth; both parameters were significantly correlated (Fig. 3B).

**Figure 3:**
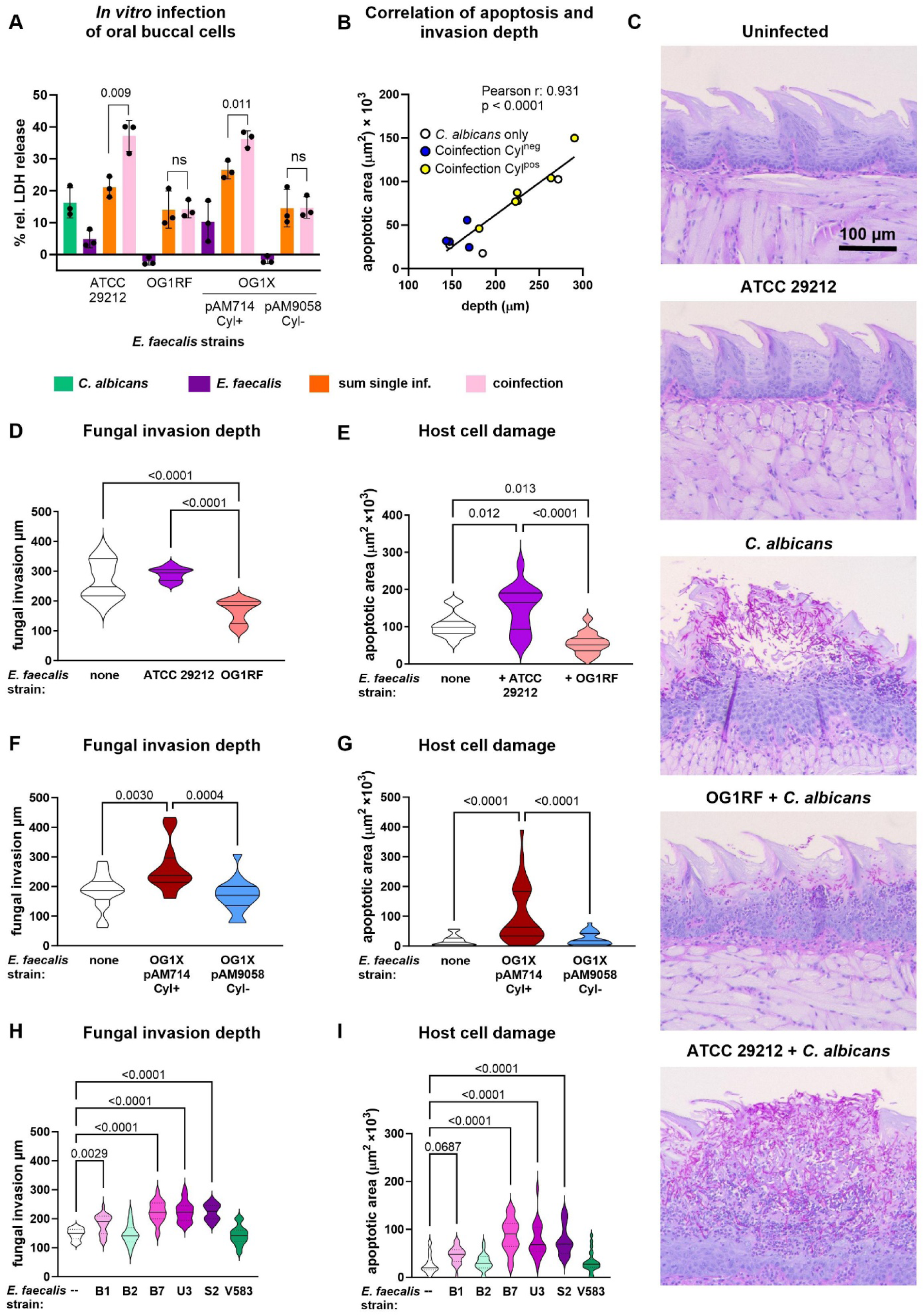
Damage of oral cells *in vitro* and murine oropharyngeal candidiasis *in vivo* are aggravated by coinfection with cytolysin-producing *E. faecalis* strains. (**A**) Human oral buccal cells (TR146) were infected with 10^5^ cells/ml *C. albicans* SC5314 for 6 h at 1 % O_2_ followed by addition of 10^5^ cells/ml *E. faecalis* and further incubation for 24 h. Damage was measured by release of host cell lactate dehydrogenase (LDH) into the supernatant, normalised to non-infected controls, and is displayed as percent damage relative to Triton X-100-lysis. Green: monoinfection with *C. albicans*; dark purple: monoinfection with *E. faecalis*; orange: sum of monoinfections; light pink: coinfection. Bars represent mean and SD, n = 3. The sum of LDH release/apoptosis caused by bacterial and fungal monoinfection was compared to coinfection using two-tailed unpaired Student’s t-tests. Significant values are indicated by p-values. ns: not significant (p > 0.05). (**B-I**) Six to seven weeks old male corticosteroid-treated C57BL/6J mice were sublingually inoculated with *C. albicans* SC5314 for 75 min. Six hours later *E. faecalis* was added to the drinking water for 24 h (2 × 10^7^ CFUs in 50 ml). (**B**) Correlation of apoptotic area and invasion depth. Mean values of all experimental groups (C-H) were analysed by Pearson correlation. (**C-E**) Coinfection with *E. faecalis* ATCC 29212 and OG1RF, respectively, n=5 mice per group. (**C**) Representative histographs of PAS-stained tongue sections, scale bar 100 µm. For each animal (n=5 mice per group), three non-consecutive sections per tongue (sections from different parts of the tongue) were randomly selected and stained. (**D, F, H**) Invasion depth was measured for each lesion visible on histological sections. Three non-consecutive section per mouse were analysed. (**E, G, I**) Size of apoptotic area was measured for each lesion visible on histological sections. Three non-consecutive section per mouse were analysed. (**D, E**) 5 mice/group. (**F, G**) Coinfection with *E. faecalis* OG1X pAM714 (cytolysin positive: Cyl+) and pAM9058 (cytolysin negative: Cyl-), respectively. Data from two independent experiments, n=6 mice/group in total. (**H, I**) Coinfection with different *E. faecalis* strains. Data from two independent experiments, n=5 mice/group in total. Pink/purple shades: cytolysin positive; green shades: cytolysin negative. (**D-I**) Data was analysed by 1-way ANOVA with Tukey’s multiple comparisons test to compare all groups (**D-G**) or Dunnett’s multiple comparisons test to compare coinfection with *C. albicans* only (**H, I**). P-values < 0.05 are indicated in the graphs.

Application of haemolytic *E. faecalis* alone, in the absence of *C. albicans* infection, did not cause any histological alterations (Fig. 3C). The consequences of *E. faecalis* on OPC were strain-dependent: The cytolysin-negative strain OG1RF, which was previously shown to reduce *C. albicans* virulence in a *Caenorhabditis elegans* model by production of EntV (Cruz et al., 2013; Graham et al., 2017), led to a significant reduction of the depth of fungal invasion (Fig. 3D) and areas of apoptosis (Fig. 3E). In contrast, the haemolytic strain ATCC 29212 led to more extensive lesions (Fig. 3D, E and Fig. S5A). Similarly, comparison of two *E. faecalis* OG1X derivatives showed enhanced tissue invasion and apoptosis only for the cytolysin-positive strain (Fig. 3F, G and Fig. S5B). By testing six additional *E. faecalis* strains, we confirmed that increased severity of OPC by *E. faecalis* depends on the presence of the *cyl* operon (Fig. 3H, I).

Plating of homogenized tongues tissue from uninfected and infected mice confirmed the absence of endogenous *C. albicans* strains (Fig. S6 A, B). Quantification of fungal burden showed a significant but minor increase in *C. albicans* CFU in co-infection with 2/10 strains (Figure S6); both *E. faecalis* strains were cytolysin positive. However, no increased fungal burden was observed with the other four haemolytic *E. faecalis* strains tested, three of which led to significantly increased tissue damage. Thus, increased fungal burden might contribute to the augmented severity of OPC for some but not all strains. The effect of *C. albicans* on *E. faecalis* CFU was variable; a significant positive effect was observed for *E. faecalis* ATCC 29212 and OG1RF (Fig. S6A), but not for *E. faecalis* OG1X (Fig. S6B). Importantly, within experiments bacterial burden did not differ between cytolysin-positive and -negative strains in either the presence or absence of *C. albicans* (Fig. S6A-C).

### Physical interaction with fungi leads to accumulation of bacteria on host cells

While soluble factors contributed to increased damage of coinfections, physical contact could also be involved. To test this, we separated microbes from host cells and/or each other by using cell culture inserts. Separation of both bacteria and fungi from enterocytes completely abolished host cell damage (Fig. 4A, orange bar), consistent with the requirement of adhesion and invasion for damage caused by *C. albicans* (Sundstrom et al., 2002; Zhu and Filler, 2010), and the lack of solubility of *E. faecalis* cytolysin in most liquid media (Booth et al., 1996; Todd, 1934). The presence of *E. faecalis* in the inserts did not affect host cell damage when *C. albicans* was in contact with enterocytes (Fig. 4A, green/turquoise bars). In contrast, a significant increase of damage by *E. faecalis* was observed if *C. albicans* was applied to the insert (Fig. 4A, purple bars), confirming the role of soluble factors in enterocyte damage. However, the damage in this setting, as well as the sum of the damage caused by either microbe on enterocytes when physically separated from the other, was significantly lower than that observed if both microbes were in physical contact with each other and the host cells (Fig. 4A, pink bar), indicating a significant contribution of contact-mediated mechanisms. Similarly, the presence of *S. cerevisiae* in the trans well insert significantly increased enterococcal host cell damage, but damage was lower than that caused by coinfection (Fig. 4A, right). Independent of physical contact, the presence of *C. parapsilosis* did not affect enterocyte damage (Fig. 4A, right). One possibility for the higher damage caused by coinfection is that adhesion of bacteria to fungal elements, as observed by electron microscopy (Fig. 4B), increases the contact of bacteria with host cells. To test this, we carefully removed the supernatant after infection and plated the supernatant and lysed host cell layer separately. While the majority of bacteria were found in the supernatant of monoinfections, significantly more bacteria were found on the host cell layer than in the supernatant after coinfection with *C. albicans*, reflecting a significant increase of bacteria on enterocytes (Fig. 4C). To test whether this effect is specific for *C. albicans*, we also analysed *S. cerevisiae*, which showed synergistic coinfection damage, and *C. parapsilosis*, which did not. *C. parapsilosis* had no effect on bacterial distribution (Fig. 4C), significantly more bacteria were isolated from supernatant than enterocytes. This is consistent with the failure of this fungus to induce synergistic coinfection damage. *S. cerevisiae* displayed intermediate results, bacterial numbers on enterocytes were increased (Fig. 4C), but the increase was not statistically significant. This could be due to the significantly weaker adhesion of *S. cerevisiae* to enterocytes compared to *C. albicans* (Fig. 4D). This difference was not only observed for enterocytes but also with the plastic surface of cell culture plates (Fig. 4E). Adhesion of *S. cerevisiae* and *C. parapsilosis* was comparable (Fig. 4D, E), suggesting differences in the interaction with *E. faecalis* might mediate the differences in bacterial location during coinfection observed for these species (Fig. 4C).

**Figure 4:**
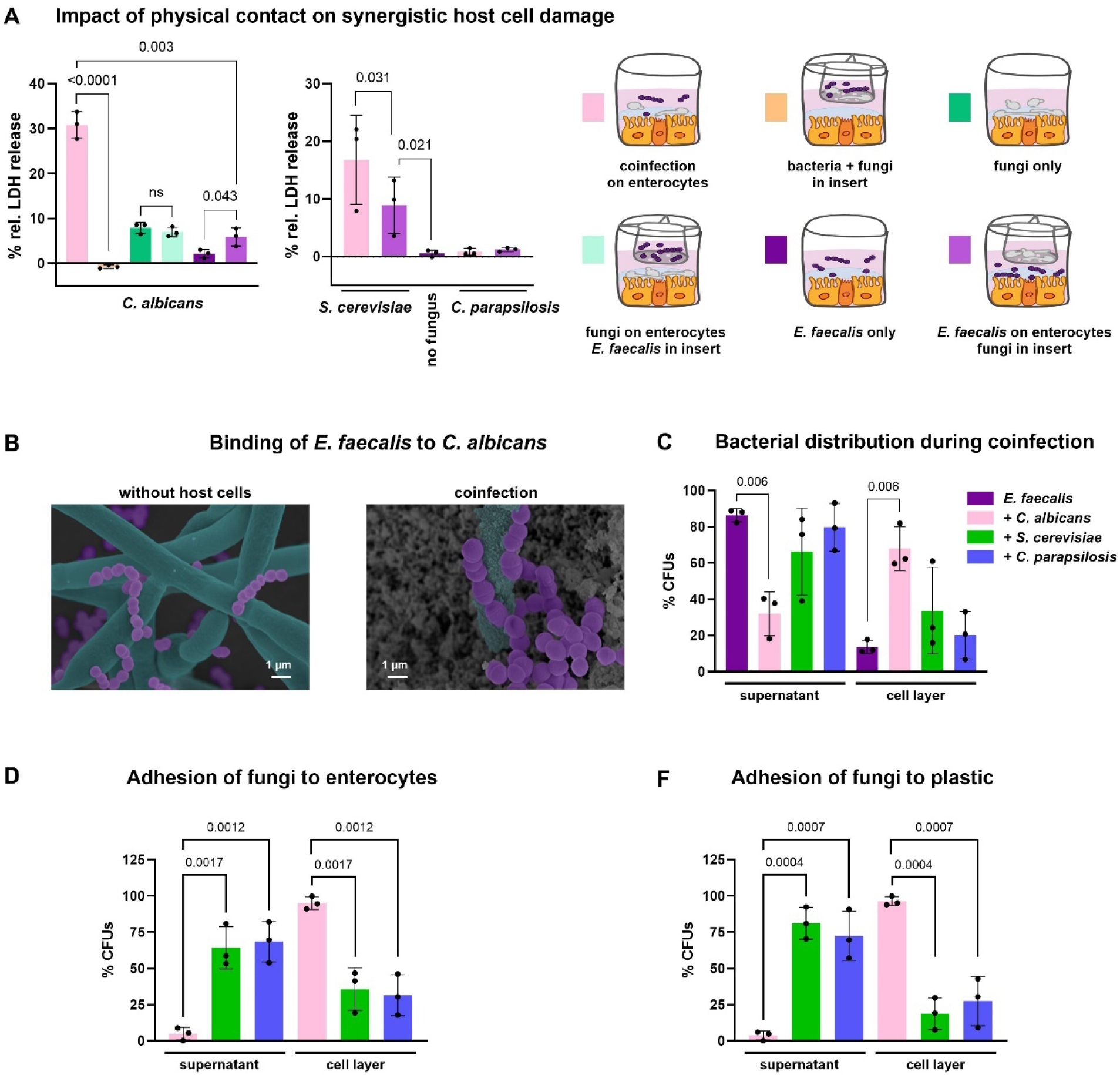
Fungal-bacterial contact leads to accumulation of bacteria at the host cell layer and downregulation of gelatinase expression. (**A**) Mono- and coinfection of enterocytes with *C. albicans* SC5314 and *E. faecalis* ATCC 29212 (left) or *E. faecalis* ATCC 29212 and *S. cerevisiae* and *C. parapsilosis*, respectively (right) using transwell inserts with 0.4 µm pores size for spatial separation. Enterocytes were infected with 10^6^ fungal cells/ml for 6 h of followed by addition of 10^5^ cells/ml *E. faecalis* and further incubation for 24 h. Damage was measured by release of host cell lactate dehydrogenase (LDH) into the supernatant, normalised to non-infected controls, and is displayed as percent damage relative to Triton X-100-lysis. The scheme below depicts the location of fungal/bacterial cells in the well or insert, respectively. Data is shown as mean with SD, n = 3. Statistical significance was analysed for *C. albicans* by two-tailored unpaired Student’s t-test with p-values indicated in the graph (ns = not significant; p > 0.05). For *S. cerevisiae* and *C. parapsilosis*, damage caused by coinfection was compared to *E. faecalis* on enterocytes with fungi in the insert by 1-way ANOVA followed by Dunnett’s multiple comparisons test; significant differences are indicated by p-value in the graph. For *S. cerevisiae*, coinfection damage and damage by *E. faecalis* on enterocytes with fungi in the insert was analysed by ratio paired t-test due to variations of coinfection damage between experiments. (**B**) Scanning electron micrographs, representative images from two independent experiments. The images were coloured manually; turquoise: fungi, fuchsia: bacteria. Without host cells: *E. faecalis* and *C. albicans* coculture in KBM for 24 h. Coinfection: Enterocyte coinfection with *E. faecalis* ATCC 29212 and *C. albicans* SC5314 (24 h). (**C**) Ratio of bacterial cells recovered from the supernatant or the host cell layer after monoinfection with *E. faecalis* ATCC 29212 or coinfection with *C. albicans* SC5314, *S. cerevisiae* or *C. parapsilosis*. Data is shown as mean with SD, n = 3, and was analysed by comparing coinfection data for supernatant and cell layer, respectively, to monoinfection by 1-way ANOVA followed with Dunnett’s multiple comparisons test. Statistical significance (p > 0.05) is indicated by p-values in the graph. (**D, E**) Ratio of fungal cells recovered from the supernatant and the host cell layer (**D**) or plastic surface (**E**) after 10^6^ fungal cells/ml were incubated for 30 h. Data is shown as mean with SD, n = 3, and was analysed by 1-way ANOVA followed with Dunnett’s multiple comparisons test comparing *C. albicans* to *S. cerevisiae* and *C. parapsilosis*, respectively. P-values are indicated in the graph.

### *Candida*-induced glucose starvation sensitizes enterocytes to bacterial damage and pore-forming toxin

To test if soluble factors produced by or changes in media composition due to the metabolic activity of *C. albicans* affect bacterial virulence, we generated *Candida*-conditioned medium (CCM) by growing fungi either alone (CCM-) or on enterocytes (CCM+) in KBM medium for 30 h. These media were then used for infection of enterocytes with *E. faecalis*. Both media led to significantly increased host cell damage during *E. faecalis* monoinfection (Fig. 5A), which was more pronounced than the effect of *C. albicans* in inserts during transwell experiments (Fig. 4A). Therefore, we hypothesized that CCM either contained specific molecules produced by *C. albicans* that cause damage or that nutrient depletion due to the metabolic activity of the fungus affected bacterial virulence. Glucose in naïve KBM (9 mM) was completely depleted after 12 h incubation of *C. albicans*, irrespective of the strain used (Fig. S7A). Of note, in the transwell assay, glucose did not diffuse freely between both compartments, and higher concentrations were observed in the well with *E. faecalis* compared to the insert with *C. albicans* (Fig. S7B). Thus, the stronger effect of CCM compared to transwell experiments could be due to differences in glucose depletion. Furthermore, while naïve KBM contains no lactate, a considerable amount (13 mM) was observed after incubation of enterocytes in medium for the duration of infection experiments. Addition of *C. albicans* resulted in three-fold lower levels of lactate, likely due to its utilization by the fungus. To determine the effect of lactate on host cell damage caused by *E. faecalis*, CCM-was spiked with lactate and used for infection of enterocytes with *E. faecalis*. Lactate addition had no significant effect on *E. faecalis* mediated damage (Fig. S7C). In contrast, glucose supplementation reduced bacterial-induced damage to the level observed with fresh media (Fig. 5A). While glucose deprivation led to higher LDH release during bacterial infection, transcription of *cylL_L_* and *cylL_S_*, encoding the two cytolysin subunits, was not altered (Fig. 5B). Similarly, no clear changes in *cylL_L_* and *cylL_S_* were observed for coincubation with *C. albicans* (Fig. S8). This suggested that cytolysin expression is not regulated by glucose. However, while both *C. albicans* and *S. cerevisiae* led to glucose depletion, *C. parapsilosis* did not (Fig. S7D), suggesting a link between glucose and synergistic damage independent of *cyl* expression. Hypothesizing that glucose affects host cell susceptibility to the toxin, we used melittin, as a second small pore forming toxin found in bee venom (Ceremuga et al., 2020). Melittin-induced host cell damage was increased in the absence of glucose (Fig. 5C), suggesting that glucose-depletion by *C. albicans* leads to enhanced effects of bacterial cytolysin by sensitizing enterocytes. Addition of melittin to enterocyte cultures infected with *C. albicans*, *S. cerevisiae* or *C. parapsilosis* resulted in synergistic damage with *C. albicans* but not *C. parapsilosis*, similar to the observations with haemolytic *E. faecalis* (Fig. 5D). In contrast to coinfection, no synergism was observed with *S. cerevisiae*. Furthermore, conditioned media of both, *C. parapsilosis* and *S. cerevisiae*, increased damage by *E. faecalis* to the extent observed for *C. albicans* (Fig. S7E). Thus, even though glucose depletion affects host cell susceptibility to melittin and bacterial damage, it does not fully explain the synergism during coinfection and the effects of conditioned media.

**Figure 5:**
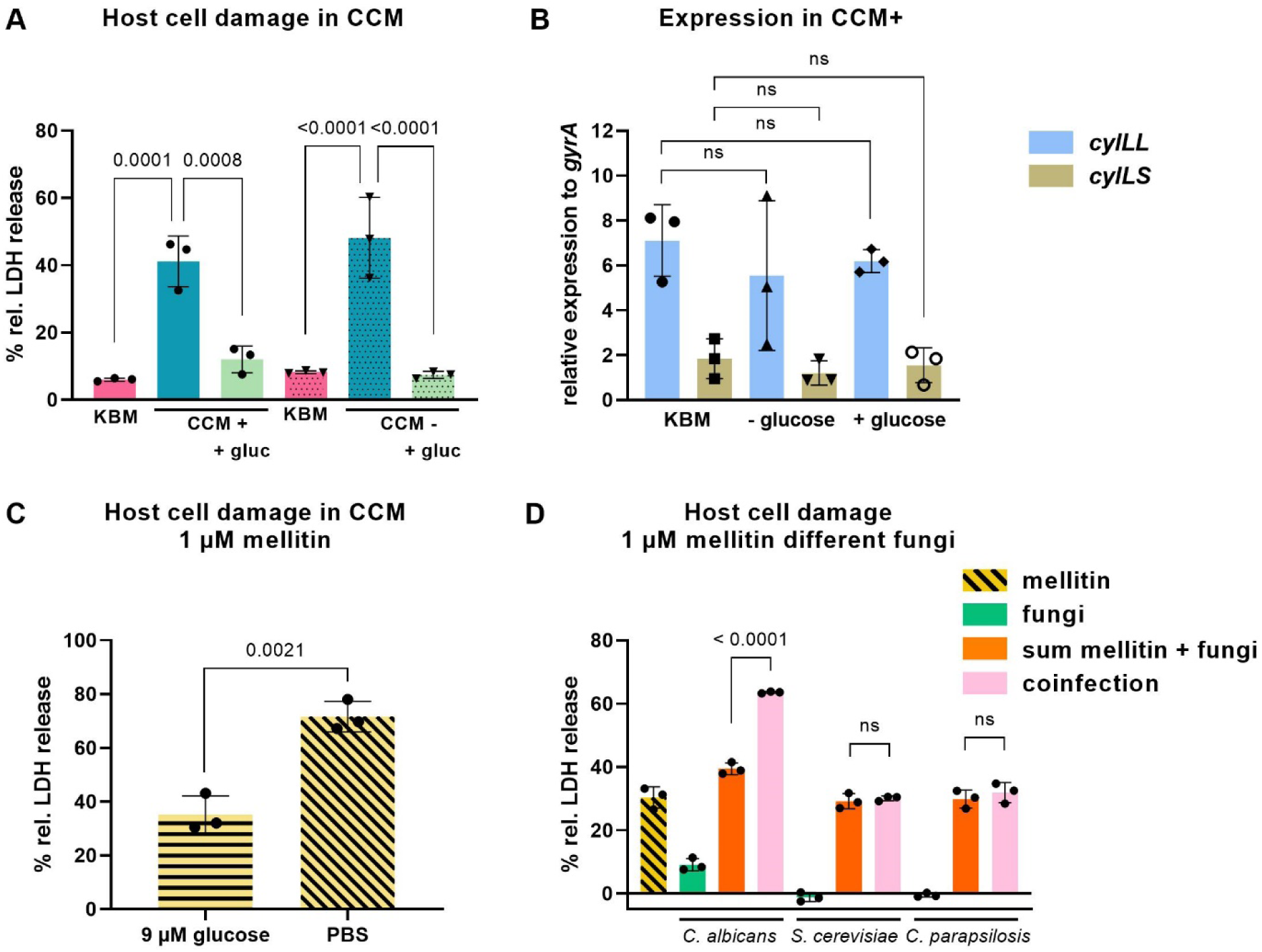
Glucose depletion does not lead to upregulation of cytolysin secretion but sensitizes host cells to damage. (**A**) *C. albicans* SC5314 conditioned media (CCM) were prepared by incubating *C. albicans* SC5314 for 30 h (1% O_2_, 5% CO_2_) in KBM with (CCM+) or without enterocytes (CCM-). The medium was sterile filtered and used for infection of enterocytes with *E. faecalis* ATCC 29212. Conditioned media were used directly or supplemented with 9 mM glucose (+gluc). Data (n=3) was analysed by 1-way ANOVA with Tukey’s multiple comparisons test to compare all groups; p-values < 0.05 of comparisons within the same type of CCM (+/-) are indicated in the graphs. (**A, C, D**) Damage was measured by release of host cell lactate dehydrogenase (LDH) into the supernatant, normalised to non-infected controls, and is displayed as percent damage relative to Triton X-100-lysis. (**B**) Expression of the large (*cylLL*) and small (*cylLS*) cytolysin subunits normalized to the housekeeping gene *gyrA* in *E. faecalis* ATCC 29212 cultured in CCM and controls for 18 h. Data (n=3) was analysed for each gene by 1-way ANOVA with Tukey’s multiple comparisons test to compare all groups. (**C**) Melittin-induced enterocytes damage in CCM+ supplemented with or without glucose (PBS as control). n = 3, data is shown as mean with SD, groups were compared with two-tailored unpaired Student’s t-test. (**D**) Effect of addition of melittin on enterocyte damage during infection with *C. albicans* SC5314, *S. cerevisiae*, or *C. parapsilosis* (n = 3 independent experiments). The sum of damage caused by the fungus and melittin separately (orange) was compared to the damage during coincubation (pink) for each fungal species with two-tailored unpaired Student’s t-test.

## Discussion

*C. albicans* and *E. faecalis* co-exist on mucosal surfaces in the human body as part of the normal microbiota. Furthermore, both thrive under dysbiosis, and the presence of *C. albicans* correlates with higher colonization levels of *E. faecalis* (Bertolini et al., 2021a; Bertolini et al., 2019; Mason et al., 2012a; Mason et al., 2012b), suggesting synergistic interactions. However, antagonistic effects on virulence mediated by the *E. faecalis* bacteriocin EntV have also been described (Cruz et al., 2013; Graham et al., 2017). This raises the question if this antagonistic interaction neutralizes the increased risk of infection usually associated with elevated fungal and bacterial colonization. We chose to address this by using an *in vitro* model that allowed us to assess the consequences of coinfection on enterocyte damage using a collection of reference and laboratory strains, as well as clinical isolates.

This approach revealed that the consequences of coinfection were highly strain-dependent. The majority of *E. faecalis* strains, including the commonly used laboratory strain OG1RF, led to a neutral outcome, in which the damage after coinfection was similar to the sum of damage inflicted by each microbe alone. A distinct set of strains, however, displayed significantly increased host cell damage after coinfection with *C. albicans*. One feature of these strains was haemolytic growth on horse blood agar plates, which is typically caused by the *E. faecalis* cytolysin (Basinger and Jackson, 1968). We confirmed the presence of the cytolysin operon in these strains, with the exception of strain ATCC 19433. This strain furthermore caused haemolysis and detectable enterocyte damage only at high oxygen concentrations, whereas coinfection synergy of the other tested strains was more pronounced at low oxygen. As cytolysin expression has been reported to be higher at low oxygen (Day et al., 2003), this together with the absence of a detectable cytolysin operon strongly suggests that ATCC 19433 carries another factor responsible for the observed haemolysis. Two strains, B3 and B8, showed no or only weak haemolytic activity on HBA and no synergistic damage *in vitro* despite presence of the complete operon in the genome. While not tested in this study, a possible explanation is a lack of cytolysin operon expression in these strains. Importantly, gain or loss of haemolytic activity by plasmid curing, plasmid transfer, and using isogenic mutants correlated with synergistic coinfection damage in different genetic strain backgrounds. This indicates that increased damage caused by coinfections can be attributed to enterococcal cytolysin.

Increased toxin production by bacteria in the presence of *C. albicans* was demonstrated for *Staphylococcus aureus* (Todd et al., 2019a; Todd et al., 2019c), but could not be confirmed in this study for *E. faecalis* on the transcriptional level. Attempts to detect and quantify the small cytolysin peptides *in vitro* experiments by unbiased proteomics were unsuccessful. Thus, while we cannot exclude that coinfection with *C. albicans* results in increased cytolysin production by *E. faecalis*, it is likely not the main or only mechanism involved.

An alternative explanation for the increased enterocyte damage upon coinfection is increased local toxin concentration on enterocytes. The importance of localized toxin secretion has recently been demonstrated for the fungal toxin candidalysin that is released in fungal invasion pockets (Mogavero et al., 2021; Swidergall et al., 2021). Consistent with this hypothesis, physical contact was necessary for full synergistic damage. Of note, cytolysin expression and the cytolytic activity of the produced toxin molecules is affected by the presence of target cells: In the absence of target cells, the toxin subunits CylL_S_ and CylL_L_ form a stable complex lacking cytolytic activity. CylL_L_ has a higher affinity to lipid membranes than CylL_S_, thereby leading to an excess of free CylL_S_ in the presence of target cells. Free CylL_S_ exceeding a threshold can bind to the membrane protein CylR1, which modulates the repressive function of CylR1 and CylR2, resulting in expression of the operon (Coburn et al., 2004; Haas et al., 2002; Roux et al., 2009). As a consequence, cytolysin-positive *E. faecalis* strains can cause haemolysis on blood agar plates, but fail to lyse erythrocytes in liquid media (Todd, 1934). In addition, quorum-sensing autoinduction depending on cell density drives cytolysin expression and adherence at the infection site promotes the toxic effects of cytolysin (Haas et al., 2002). The presence of *C. albicans* resulted in a higher number of bacterial cells in association with enterocytes, and thereby possibly enhanced cytolytic activity.

Binding to fungal cells has also been reported for *S. aureus* and oral streptococci (Diaz et al., 2012; Peters et al., 2012). While *S. aureus* binds preferably to Als3 expressed on hyphae (Peters et al., 2012), synergistic damage by *E. faecalis* also occurred with filament-deficient *C. albicans* strains and other yeast species, suggesting a different binding mechanism. The synergistic coinfection damage observed with non-filamentous fungi furthermore implies that pronounced fungal invasion and creation of invasion pockets are not necessary for increased cytolysin-mediated damage, although they did contribute to overall host cell damage. *S. cerevisae*, which led to increased damage in *E. faecalis* coinfection, also trended to increase the number of bacteria on host cells. It should be noted that binding of *S. cerevisiae* to host cells and the plastic surface was weaker than that of *C. albicans*. This possibly resulted in the loss of bacteria during washing steps, and consequently underestimation of the number of bacteria in contact with enterocytes during coinfection with *S. cerevisiae*. Furthermore, coinfection with *C. parapsilosis* had no effect on bacterial association with the host cell layer and did not result in synergistic damage. As the adhesion of *C. parapsilosis* and *S. cerevisiae* to host cells and the plastic surface was comparable, different factors might mediate binding to surfaces and interactions with *E. faecalis*.

Taken together, our results suggest that physical interactions between fungi and bacteria, resulting in an increased concentration of bacteria on host cells enhances cytolysin-mediated damage. This would be consistent with the enhancing effects that target cells have on cytolysin regulation (Coburn et al., 2004; Haas et al., 2002; Roux et al., 2009), haemolysis on solid but not in liquid media (Todd, 1934), and the lack of damage enhancement if bacteria were located on the insert during transwell experiments.

While physical interaction played a prominent role for coinfection damage *in vitro*, culture-conditioned media also had an effect on enterococcal host cell damage. This implied that consumption of nutrients or release of small molecules by *C. albicans* could affect bacterial virulence. This might also explain why synergistic damage *by C. albicans* only occurred if enterocytes were infected with *C. albicans* first, and why the synergistic effect was more pronounced with increasing preincubation time and fungal cell numbers: Ongoing fungal metabolic activity would lead to increased nutrient depletion or conversion over time. Production of small molecules, including secreted metabolites, might be more pronounced in the absence of bacterial nutrient competition. Here, we focused on glucose consumption by *C. albicans* as an activity that would lead to depletion of nutrients available to the bacteria. While glucose depletion increased host cell damage by *E. faecalis*, this was not attributable to increased cytolysin production. Furthermore, the use of KBM medium with a high buffering capacity prevented media acidification during the experiments. Therefore, we considered effects of glucose on host cells. Because recombinant cytolysin is not available, we used melittin as a model pore-forming toxin, and observed that damage by melittin was reduced by glucose supplementation. A protective effect of luminal glucose was also observed for LPS-, and *Giardia*-induced enterocyte apoptosis *in vitro* (Yu et al., 2005; Yu et al., 2008), and ischemia/reperfusion-induced enterocyte damage and barrier dysfunction *in vivo* (Huang et al., 2011). While the mechanisms by which glucose reduces toxin-mediated damage remain unclear, the negative effects of fungal glucose depletion have been demonstrated for *C. albicans* and macrophages. Infected macrophages increase glycolysis and rely on glucose for survival. *C. albicans* rapid glucose uptake depletes glucose available for the macrophage, thereby causing macrophage death (Tucey et al., 2018). This illustrates that nutrient depletion by microbes can affect host cell susceptibility. Interestingly, though, the addition of melittin to enterocytes infected with *C. albicans*, *S. cerevisiae* or *C. parapsilosis* resulted in synergistic damage with *C. albicans* only, whereas coinfection of *E. faecalis* also resulted in synergistic damage with *S. cerevisiae*. This indicates differences between the toxins, and that results obtained with melittin cannot be directly translated to enterococcal cytolysin. Furthermore, infection of enterocytes with *E. faecalis* in conditioned media from all three fungal species resulted in increased host cell damage. This cannot be attributed to glucose depletion, and for *C. parapsilosis* is in stark contrast to the lack of synergistic damage in coinfection experiments. It seems possible that various soluble factors influence host cell susceptibility either positively or negatively, and that the ultimate outcome depends on the quality and quantity of different factors.

Cytolysin-dependent synergistic damage *in vitro* was not restricted to enterocytes but also occurred with oral epithelial cells. Since intestinal colonization of mice with neither *C. albicans* nor *E. faecalis* results in detectable epithelial damage (Garsin et al., 2014; Garsin and Lorenz, 2013; Shao et al., 2022), we decided to use a murine oropharyngeal candidiasis (OPC) model in which mucosal damage can be more easily measured to test if cytolysin-dependent synergistic damage occurs *in vivo*. Furthermore, physical interaction between *C. albicans* and *E. faecalis* has been observed during murine OPC (Bertolini et al., 2019). The results confirmed increased host damage by coinfection with *C. albicans* and cytolysin-producing *E. faecalis*. Additionally, the cytolysin-negative strain OG1RF led to reduced fungal invasion depth. This likely reflects EntV-mediated reduction of filamentation and virulence, which was described in other infection models (Cruz et al., 2013; Graham et al., 2017), but was not evident in *in vitro* enterocyte infection. While it appears likely that the physical interactions between *C. albicans* and *E. faecalis in vitro* also affect *in vivo* coinfection, the role of metabolism and soluble factors might differ. Regarding glucose availability, for example, it should be noted that normal blood glucose levels (70 – 100 mg/dl) are only slightly lower than the amount of glucose in cell culture media, and remain relatively constant over time, providing continuous supply for tissues. However, changes in local perfusion due to inflammation might create microenvironments in which glucose concentrations are lower independent to microbial activities, or increased blood flow might compensate for increased glucose consumption. It therefore remains unclear if and to which extend *C. albicans*’ metabolic activity affects glucose concentrations *in vivo*.

In summary, this study demonstrates that the outcome of *C. albicans* – *E. faecalis* coinfection depends on strain-specific bacterial properties, and we identified cytolysin as the factor aggravating host damage *in vitro* and *in vivo*. Therefore, species identification alone might not be sufficient to predict the risk of infections in patients with concurrent bacterial and fungal overgrowth. Furthermore, both physical interactions between the microbes and metabolic activity altering the environment and affecting host cell resilience contribute to increased damage during coinfection.

## Materials and Methods

### Strains and culture conditions

All strains used in this study are listed in Table S4 and S5. Strains were stored as glycerol stocks at -80 °C and streaked on solid media prior to experiments. Fungal overnight cultures were prepared by inoculating 10 ml yeast extract peptone dextrose (YPD; 2% glucose, 2% peptone and 1% yeast extract) with a single colony and incubation at 30 °C and 180 rpm. Bacteria were streaked on Columbia CNA Agar (HiMedia) or Brain Heart Infusion (BHI) Agar (Carl Roth) containing 5% horse blood (Thermo Scientific). For overnight cultures, colonies were transferred into 5 ml liquid Todd-Hewitt-Broth medium (THB) (Carl Roth) and incubated at room temperature (RT) at 180 rpm. For experiments, overnight cultures were inoculated to OD_600_ 0.1 in fresh THB and grown for approximately 2 h at 37 °C and 180 rpm till they reached an OD_600_ of approximately 1. The bacterial OD_600_-CFU-correlation was determined by plating serial dilutions of such cultures on lysogeny broth agar (LB; 1% tryptone, 0.5% yeast extract, 1% sodium chloride) and quantification of colony forming units (CFUs) after 24 h incubation at 37 °C. Heat inactivation was achieved by incubating cultures diluted to the desired concentrations for 1 h at 65 °C in a water bath. Killing was confirmed by plating on THB (bacteria) or YPD (fungi) agar plates.

### *In vitro* infection models

For enterocyte infection a 7:3 mixture of the two human cell lines C2BBe1 (brush-border expressing enterocytes, ATCC CRL2102) and mucus producing HT29-MTX-E12 (mucus producing, European Collection of Authenticated Cell Cultures Cat. No.: 12040401) cells were grown at 37 °C with 5% CO_2_ in Dulbecco’s Modified Eagle Medium (DMEM, Gibco) supplemented with 10% v/v fetal calf serum (FCS) (Bio&Sell), 0.01 mg/ml Holo-Transferrin (Sigma), and 1% non-essential amino acids (Glibco) (Hilgendorf et al., 2000). Cells were differentiated for 14 days with medium exchanges twice a week. TR146 cells (human oral buccal cells, European Collection of Authenticated Cell Cultures Cat. No.: 10032305) were grown in DMEM/F12 (Gibco) with 10% FCS for 48 h. Fungal overnight cultures were washed twice with phosphate buffered saline (PBS), counted in a Neubauer Chamber (Hecht Assistent), and set to 2×10^7^ cells per ml in PBS. Bacteria were washed twice with PBS and diluted to a cell density of 2×10^6^ per ml based on the CFU-OD_600_-correlation. Host cells (mixed enterocytes or oral buccal cells) were washed once with PBS and KBM (Gold Keratinocyte Medium; Lonza Walkersville) or conditioned KBM was added. KBM was used instead of the respective cell culture maintenance medium due to its higher buffering capacity to prevent media acidification by bacteria during infection. Then, host cells were infected with 1×10^6^ fungal cells per ml (MOI = 1; e. g. 10 µl of 2×10^7^ cells stock solution in 200 µl for the 96-well plate format; 50 µl for 24-well plates; 250 µl for 6-well plates) unless stated otherwise, and incubated for 6 h at 37 °C, with 5% CO_2_ and either 21%, 10%, or 1% O_2_. After 6 h (or earlier, if indicated) 1×10^5^/ml enterococci were added. After additional 24 h of incubation supernatants were removed, and samples were processed as described below. Due to the fast growth of the bacteria, bacterial CFU exceeded fungal CFU at the end of the experiment (Fig. 1S). Trans-well experiments were performed as described above in 24 well plates using inserts with a 0.4 µm pore size (Merck). In order to assess host cell susceptibility to small pore forming toxins, 1 µM of melittin (Sigma) was added to the host cells as described in (Ceremuga et al., 2020).

### Quantification of host cell damage

Host cell damage was quantified by measuring lactate dehydrogenase (LDH) release into the supernatant using the Cytotoxicity Detection Kit (Roche) according to the manufacturer’s instructions. For each experiment, supernatants from uninfected cells and uninfected cells lysed with 0.25% Triton X-100 (Sigma) served as negative and positive controls, respectively. Infection-mediated cell damage was calculated by subtracting the non-infected control values from all other samples relative to the high control.

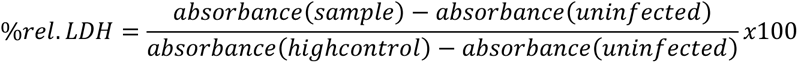

### Quantification of host cell apoptosis

To measure apoptosis, 5% DMSO (Carl Roth) was added as positive control to non-infected host cells for 4h. Apoptotic cells were measured by using FAM-DEVD-OPH *in vitro* Apoptosis Detection Reagent (ICT-6356) according to the manufacturer’s protocol. Briefly, 700 μl of supernatant was removed and 10 μl staining solution was added to each well. After 1-hour incubation at room temperature, cells were scrapped off followed by washing once with PBS. Cells were resuspended in 300 µl of PBS and mixed gently to disrupt clumping. Apoptotic cells were quantified on a fluorescence plate reader (Tecan infinite M200 pro) at Ex/Em = 490/525 nm.

### Quantification of total microbial burden during host cell infection

To quantify microbial burden, 10 µl supernatants were removed at the indicated time points for host cell damage quantification the wells were then treated with 1 mg/ml Zymolyase 20T (amsbio) for 30 min at 37 °C. The well content was scraped off, removed from the well, and further incubated for 30 min to separate fungal hyphae. The samples were serially diluted and plated on YPD agar containing 50 µg/ml gentamicin and 100 µg/ml doxycycline (for quantification of fungi) and LB agar containing 50 µg/ml nystatin (for quantification of bacteria). Colonies were counted after 24 h incubation at 37 °C.

### Quantification of bacterial cell numbers in infection supernatants and on host cells

Single and coinfections were carried out as described above. At the end of infection, the supernatant was used to carefully wash off loose bacteria by pipetting up and down once before it was removed from the cell layer. The remaining cell layer was then washed once and resuspended in PBS by rigorous pipetting. Both supernatant and cell fraction were serially diluted and plated on LB agar to quantify bacterial CFUs.

### Generation of *C. albicans* conditioned media

Enterocytes or empty wells containing KBM only were inoculated as described above with 1×10^6^ cells/ml *C. albicans* in 30 ml and incubated for 30 h at 37 °C with 5% CO2 and 1% O_2_. The supernatant was removed, filtered through a 2 µm Filtropur sterile filter (Sarstedt), and stored at -80 °C. Conditioned medium was used either directly or spiked with glucose (Carl Roth) or L-(+)-lactate (Sigma) to a final concentration of 9 mM or 13 mM, respectively.

### Quantification of glucose and lactate in cell culture supernatants

Host cells were either uninfected or infected with ether 1×10^6^ cells/ml fungi, 1×10^5^ cells/ml *E. faecalis*, or both. Over a course of 10 h each hour one droplet of supernatant was taken and the glucose (mmol/l) measured using a Contour blood sugar meter (Ascensia Diabetes Care, Germany). Lactate in cell culture media or culture supernatants was quantified by the Institute of Clinical Chemistry and Laboratory Diagnostics, Jena University Hospital, Jena, Germany, using routine methods.

### *E. faecalis* plasmid curing and conjugation

Plasmid curing was performed as described by McHugh and Swartz (McHugh and Swartz, 1977). Briefly enterococcal overnight cultures were diluted to 1×10^5^ cells/ml in 10 ml fresh THB medium containing 8 µg/ml novobiocin (Sigma) and incubated at 37 °C without shaking for 18 h. Afterwards the culture was serially diluted and plated on CNA Colombia Agar (HiMedia) supplemented with 5% horse blood (Thermo Scientific) overnight to determine haemolytic activity of colonies. Non-haemolytic colonies were subcultured twice on CNA Colombia Agar (HiMedia) supplemented with 5% horse blood to confirm absence of haemolysis before long-term storage in glycerol at -80°C. Strains that underwent the curing procedure but were still haemolytic were used as controls in infection experiments. To facilitate plasmid transfer by conjugation, filter mating was performed as described by Haug *et al*. (Haug et al., 2010). In short, donor (B1) and recipient (OG1RF) overnight cultures were mixed 1:3 and 100 µl of cells were dropped onto a sterile 0.45 µm nitrocellulose filter (GVS North America) placed on a nonselective THB agar plate. The plates were incubated overnight, the filter removed, and washed with 5 ml sterile PBS in a 50 ml centrifuge tube by vigorous vortexing. The suspension was then serially diluted and plated on CNA Colombia Agar supplemented with 5% horse blood and 50 µg/ml rifampicin (Sigma) overnight to select for the recipient strain OG1RF (rifampicin-resistant). Haemolytic colonies were subcultured twice on CNA Colombia Agar supplemented with 5% horse blood to confirm acquisition of haemolytic activity before long-term storage in glycerol at -80°C. Non-haemolytic colonies from the same experiments were used as negative control strains.

### Bacterial DNA Isolation

Genomic DNA from 10 ml Enterococci overnight cultures was isolated using the Qiagen Genomic-tip 100/G (Qiagen) Kit according to the manual for gram positive bacteria. Genomic DNA for smaller volumes was isolated with the kit Qiagen DNeasy Blood & Tissue according to the manual section for gram positive bacteria.

### Whole Genome Sequencing of enterococcal DNA

DNA was quantified using the Qubit High Sensitivity DNA kit (ThermoFisher). DNA was fragmented using Covaris microTUBE AFA Fiber tubes (Covaris). Fragmented DNA was used to prepare sequencing libraries using the TruSeq Nano kit (Illumina) to obtain 550 bp insert size libraries. Quality of sequencing libraries was assessed using the QIAxcel Advanced System (Qiagen). Quantity was determined using Qubit High Sensitivity DNA kit (Thermo Fisher). Sequencing libraries were sequenced on Illumina Nextseq creating 300bp paired-end reads. Sequencing coverage was 100-fold.

The raw binary base call files produced from the Illumina Nextseq were converted to fastq using Illumina’s bcl2fastq conversion software (v2.20.0.422). Before analysis, general quality readings were collected using FastQC (0.11.4 34) (Andrews, 2010), for quality control of data generated on Illumina platforms. Reference sequence data for cytolysin operon was retrieved from the Kyoto Encyclopedia of Genes and Genomes (KEGG - www.genome.jp) entry for the Enterococcus faecalis ATCC 29212 (T03320) p1plasmid (genes DR75_2951 - DR75_2958). These sequences were then compared to sequence data using the BBduk (Decontamination Using Kmers) tool from the BBMap suite (38.90) (Bushnell, 2014.), while simultaneously trimmed of lower quality regions, using the flags/options: quality trim reverse/forward ends (qtrim) = rl, trim read length equal to zero modulo 5 to correct for erroneous Illumina extra sequencer bases (ftm) = 5, trim regions with average quality below (trimq) = 10, kmer length (k) = 31, allowing one mismatch (hdist) = 1. Each cytolysin gene sequence was probed separately and the resulting kmer contamination statistics files were then used to determine its presence/absence in every strain. The sequence data was then also probed using the sequences of other known *Enterococcus* virulence factors retrieved from the Virulence Factors of Pathogenic Bacteria database (http://www.mgc.ac.cn/VFs/main.htm), to determine their presence in the strains.

Additional sequencing was conducted using the Nanopore sequencing technology (ONT). The ZymoBIOMICS DNA Miniprep Kit (Zymobiomics) was used to isolate bacterial DNA for long read sequencing. DNA quantification steps were performed using the dsDNA HS assay for Qubit (Invitrogen, US). DNA of the samples was size selected by clean up with 0.45 × volume of Ampure XP buffer (Beckman Coulter, Brea, CA, USA) and afterwards eluted in 50 μl EB buffer (Qiagen, Hilden, Germany). The library was prepared from 1 µg DNA using the SQK-LSK114 kit (Oxford Nanopore Technologies, Oxford, UK) according to the manufacturer’s protocol. Reads of the samples were basecalled using *Dorado* (v7.3.11) (Oxford Nanopore Technologies) using the high super accuracy basecalling model. The basecalled reads were assembled and polished using *Flye* (v2.9.3), *Medaka* (v1.11.3), and *Racon* (v1.4.20).

### PCR of enterococcal genomic DNA

Polymerase chain reaction with Phusion High-Fidelity DNA-Polymerase (ThermoFisher) was performed as stated in the manual with the following settings. 35 cycles, denaturation 10 s, annealing at 55 °C 30 s, elongation 45 s, final extension 10 min. Genomic DNA was diluted 1:100 and used as 1 µl. Final reaction volume 20 µl. All primers used to amplify cytolysin genes are shown in Table S4.

### Quantitative real-time PCR

To measure cytolysin transcription single- and coinfections were carried out as described above for 18 h post enterococcal infection. To assess if contact to *C. albicans* alone is sufficient to trigger cytolysin transcription single- and cocultivations were carried out using KBM medium mixed with lysed frozen enterocytes (10^6^ cells/ml) to allow optimal enterococcal growth. Microbes were scraped off, washed once with ice cold distilled water to remove enterocytes, collected into ceramic bead tubes (Genaxxon bioscience), and stored at -80 °C till further use. Frozen cell pellets were resuspended in 1 ml TRI Reagent (RiboPure™ RNA Purification Kit, #AM1924, Invitrogen). Cells were mechanically disrupted on a FastPrep-24 cell-homogenizer (MP Biomedicals) with two 30 s runs at 6.0 m/s and with 5 min on ice incubation between each run. Samples were centrifuged at 13,000 rpm for 10 min at 4°C and further processed using the RiboPure RNA Purification Kit (#AM1924, Invitrogen) according to the manufacturer’s protocol. The purified RNA was treated with TURBO DNase (#AM2238, Ambion) by adding 0.1 volume 10× TURBO DNase buffer and 1 µl TURBO DNase to each sample. DNase activity was inactivated by adding 0.1 volume DNase Inactivation Reagent. The quality and quantity of the RNA was determined by utilizing the Bioanalyzer device (Agilent) and Nanodrop (ThermoFisher), respectively. Subsequently, 1500 µg RNA was transcribed into cDNA by using the SuperScript IV Reverse Transcriptase (#18090010, Invitrogen) according to the manufacturer’s protocol. The following qRT-PCR was conducted in biological and technical triplicates with the Brillant II SYBR Green QPCR Master Mix (#600828, Agilent) in the Stratagene Mx3005 device (Agilent). In brief, 1:10 dilutions of each cDNA sample were used. The primer efficiency was determined with genomic DNA of the *E. faecalis* strain ATCC 29212 in 1:10 dilutions from 100 µg/ml to 0.1 µg/ml. The respective gene expression was calculated with the Δct method relative to the housekeeping gene *gyrA*. All primers are listed in Table S4

### Mouse model of oropharyngeal candidiasis

Six to seven weeks old male C57BL/6J mice were purchased from Jackson laboratories (USA) and housed for 1 week in a pathogen free facility prior to infection. The mice were randomly assigned to the infection groups. No antibiotics were used for infections with the other strains. Oral *C. albicans* infection was induced as described previously (Millet et al., 2020; Solis and Filler, 2012; Swidergall et al., 2018). Briefly, on day -1 and +1 post oral fungal infection mice were given a subcutaneous injection of 150 mg/kg cortisone acetate (Sigma-Aldrich). For inoculation, the animals were sedated, and a swab saturated with 1 × 10^6^ *C. albicans* cells SC5314 (Fonzi and Irwin, 1993) was placed sublingually for 75 min. Six hours post *C. albicans* infection *E. faecalis* was added to the drinking water for 24 h (2 × 10^7^ CFUs in 50 ml). This approach was chosen to resemble the *in vitro* experiments, in which synergistic damage was most pronounced if fungi were allowed to adhere to host cells before enterococci were added. Water consumption was monitored in some experiments and found to be 5 ml ± 0.4 ml/ mouse and 24 h. Thus, the effective infectious dose for *E. faecalis* is estimated to be 2 × 10^7^ CFU/mouse.

Preliminary experiments showed that *E. faecalis* OG1X + pAM714 and OG1X + pAM9058, respectively, could not be retrieved from the tongues of mice infected by the protocol described above. We considered two possible reasons: Loss of the plasmids due to the lack of selective pressure, and reduced fitness in competition with other bacteria, especially endogenous enterococci. The later hypothesis was based on the observation that these strains grew more slowly *in vitro* than other *E. faecalis* strains used. Therefore, mice to be infected with these strains were started on antibiotics (gentamicin 15 μg/mL; erythromycin 50 μg/mL) in the drinking water at day -2, and antibiotic supplementation was continued throughout the experiment. Erythromycin was used to ensure plasmid maintenance, gentamicin to suppress endogenous bacteria (OG1X is gentamicin resistant). The mice were euthanized by pentobarbital overdose followed by cervical dislocation two days after *C. albicans* infection. For CFU enumeration the tongues were harvested, cut in half, weighed, homogenized and quantitatively cultured. Sabouraud dextrose agar (SDA) containing chloramphenicol (50 μg/ml) was used for quantification of *C. albicans*. BHI Agar with 5% horse blood, gentamicin (15 μg/ml)/erythromycin (50 μg/ml), and fluconazole (250 μg/mL; Sigma-Aldrich) was used for selective culture of *E. faecalis* OG1X pAM714 and OG1X pAM9058 based on the resistance marker located on the respective plasmid. The presence or absence of haemolysis confirmed that the isolated colonies showed the anticipated phenotype. Selection by antibiotics was not possible for the other *E. faecalis* strains. Therefore, *Enterococcus* ChromoSelect Agar (Sigma-Aldrich) supplemented with fluconazole (250μg/mL) was used as selective medium for CFU quantification for all other strains. The presence/absence of haemolytic activity of colonies growing on *Enterococcus* ChromoSelect Agar was confirmed by plating randomly selected colonies on BHI Agar with 5% horse blood.

For histology and immunhistochemistry, tongues were cut in half, fixed in zinc buffered formalin, and embedded in paraffin. Three non-consecutive sections per tongue (sections from different parts of the tongue) were randomly selected and stained with Periodic Acid Schiff (PAS) were imaged by light microscopy, and the depth of individual fungal lesions relative to surface of the tongue (Solis et al., 2017) were determined using PROGRES GRYPHAX® software version 1.1.8.153 (Jenoptik). Apoptotic cells were stained by immunohistochemistry using the ApopTag *in situ* apoptosis detection kit (EMD Millipore) as described previously (Soliman et al., 2021). Briefly, three non-consecutive paraffin-embedded sections per tongue were randomly selected, rehydrated in Histo-Clear II (National Diagnostics) and alcohols followed by washing with PBS. The sections were pre-treated with 20 μg/mL Proteinase K (Ambion) in PBS 15 min at RT. Endogenous peroxidases were blocked by incubation of the slides for 15 min in 3% hydrogen peroxide. Sections were incubated with equilibration buffer (EMD Millipore) for 30 sec at RT, followed by terminal deoxynucleotidyl transferase (TdT; EMD Millipore) at 37°C for 1 h. Sections were further exposed to anti-Digoxignenin for 30 min at RT, and the positive reaction was visualized with 3-diaminobenzidine substrate (Thermo Scientific). After counterstaining the specimens with 0.5% methyl green (Sigma), they were imaged by bright field microscopy. For quantification, apoptotic areas were quantified using PROGRES GRYPHAX^®^ software (Jenoptik). Samples were blinded before evaluation.

### Ethics statement

All animal work was approved by the Institutional Animal Care and Use Committee (IACUC) of the Lundquist Institute at Harbor-UCLA Medical Center (protocol number 30927-01R).

### Statistical analysis

Statistical analyses were performed using Graphpad Prism version 9.2.0. The type of statistical analysis used is indicated for each data set in the figure legend.

## Acknowledgments

We want to thank Katja Schubert for outstanding technical support. We further want to thank Steffen Höring, Oliwia Makarewicz, Danielle Garsin, and Michael Lorenz for providing us with bacterial isolates. Furthermore, we want to thank the Department of Microbial Pathogenicity Mechanisms (HKI) especially Selene Mogavero and Sascha Brunke for supplying us with bacterial isolates and human host cell lines as well as for encouraging discussions and support. We are grateful to the members of the research group Host Fungal Interfaces for the valuable input and discussions.

## Funding

This project was supported by the Bundesministerium für Bildung und Forschung (BMBF) via the Center for Sepsis Control and Care, grant 01EO1002, as part of the projects “CanBac: Candida and Bacteria” (IDJ) and “GuLiver” (ASM) the TRR 124 FungiNet “Pathogenic fungi and their human host: Networks of Interaction,” Deutsche Forschungsgemeinschaft (DFG, German Research Foundation), project number 210879364, projects C5 (IDJ) and C2 (SV)

NIH grant R00DE026856 (MS)

NIH grant R01DE031382 (MS)

American Association of Immunologists Careers in Immunology Fellowship (MS)

Bundesministerium für Bildung und Forschung (BMBF) via the funding program Photonics Research Germany, Leibniz Center for Photonics in Infection Research (LPI), subproject LPI-BT4, contract number 113N15716 (ASM) and subproject LPI-BT2, contract number 13N15705 (IDJ)

the Germanýs Excellence Strategy, DFG, EXC 2051 – Project-ID 390713860 (IDJ)

## Author contributions

Conceptualization: IDJ; methodology: MK, MJN, MS, IDJ; investigation: MK, MJN, EKC, AC, KAM, MM, MS, NM, PB, AL, MV, SN, FH, ASM; visualization: MK, IDH, MS; supervision: IDJ, BL, OM, MS, KAK, SV; writing—original draft: MK, IDJ; writing—review & editing: all authors.

## Data and materials availability

All data needed to evaluate the conclusions in the paper are present in the paper and/or the Supplementary Materials.

