## Supplemental figures and tables for "Synergistic cross-kingdom host cell damage between *Candida albicans* and *Enterococcus faecalis*"

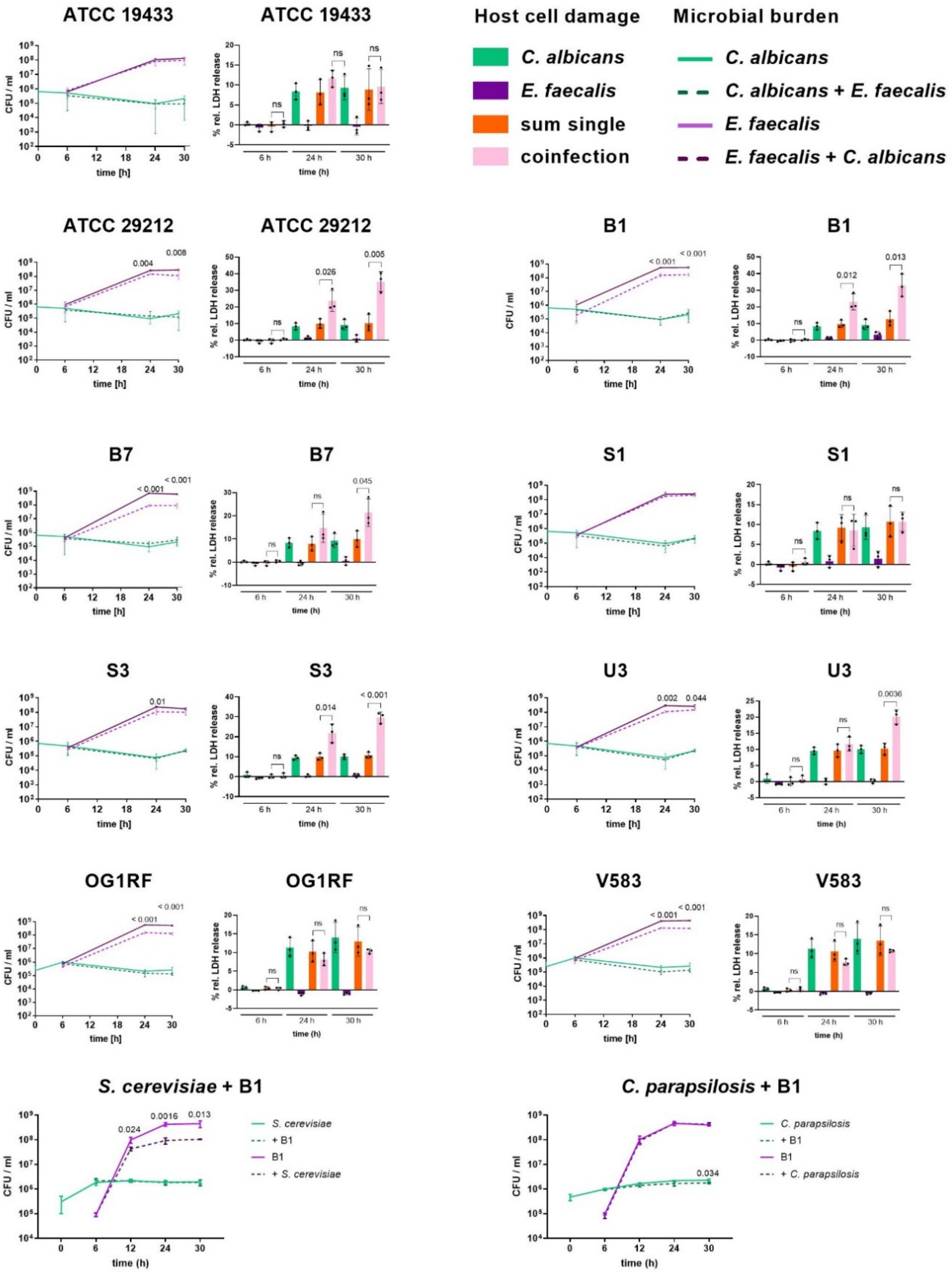

**Fig. S1: Increased damage upon coinfection is not associated with increased microbial proliferation**

Enterocytes were infected with $10^6$ cells/ml *C. albicans* SC5314 for 6 h at 1 % $O_2$ followed by addition of $10^5$ cells/ml *E. faecalis*. At the indicated time points 10 µl supernatant were removed to quantify LDH; the remaining supernatant and cell layer were then treated with Zymolyase (1 mg/ml) for 1 h at 37 °C. Cells were resuspended in the supernatant by scratching and vigorous pipetting; serially dilutions of the samples were plated on YPD (50 µg/ml Gentamycin and 100 µg/ml Doxycycline) and LB (50 µg/ml Nystatin). Damage was measured by release of host cell lactate dehydrogenase (LDH) into the supernatant, normalised to non-infected controls, and is displayed as percent damage relative to Triton X-100-lysis. Data is shown as mean with SD, n = 3. Statistical significance was analysed by two-tailored unpaired Student's t-test; p-values $< 0.05$ are indicated in the graphs. ns: not significant ($p > 0.05$).

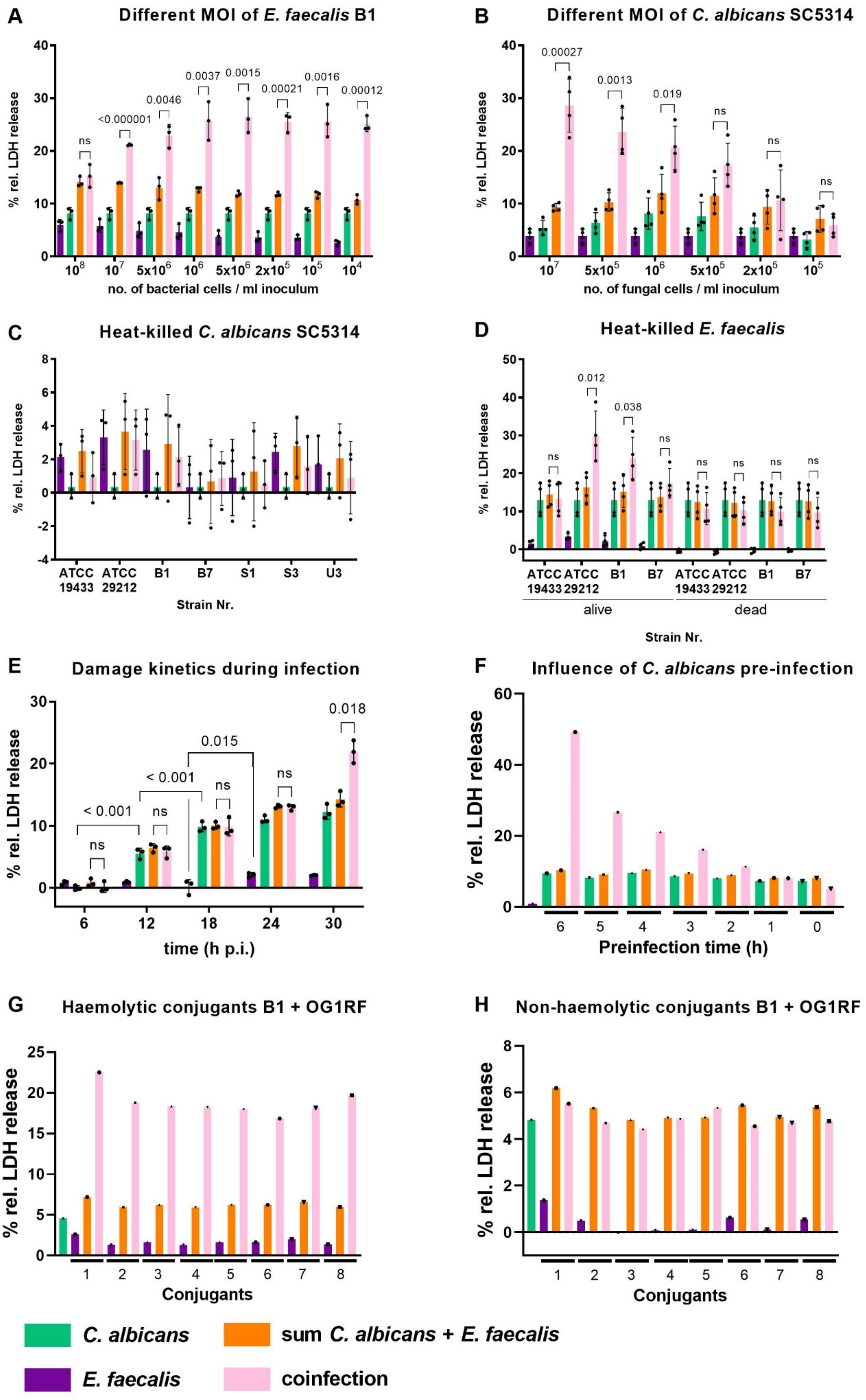
A
Different MOI of E. faecalis B1
% rel. LDH release
no. of bacterial cells / ml inoculum
ns
<0.000001
0.0046
0.0037
0.0015
0.00021
0.0016
0.00012
B
Different MOI of C. albicans SC5314
% rel. LDH release
no. of fungal cells / ml inoculum
0.00027
0.0013
0.019
ns
C
Heat-killed C. albicans SC5314
% rel. LDH release
ATCC 19433
ATCC 29212
B1
B7
S1
S3
U3
Strain Nr.
D
Heat-killed E. faecalis
% rel. LDH release
0.012
0.038
ns
alive
dead
Strain Nr.
E
Damage kinetics during infection
% rel. LDH release
time (h p.i.)
< 0.001
0.015
0.018
ns
F
Influence of C. albicans pre-infection
% rel. LDH release
Preinfection time (h)
G
Haemolytic conjugants B1 + OG1RF
% rel. LDH release
Conjugants
H
Non-haemolytic conjugants B1 + OG1RF
% rel. LDH release
Conjugants
C. albicans
sum C. albicans + E. faecalis
E. faecalis
coinfection

**Fig. S2. Synergy depends on fungal but not bacterial cell number and requires viable microbes**

Enterocytes were infected with $10^6$ cells/ml *C. albicans* SC5314 (unless indicated otherwise) at 1 % $O_2$ for 6 h followed by addition of *E. faecalis*. Damage was measured by release of host cell lactate dehydrogenase (LDH) into the supernatant, normalised to non-infected controls, and is displayed as percent damage relative to Triton X-100-lysis. **A:** Coinfection with $10^6$ cells/ml *C. albicans* and different cell numbers of *E. faecalis* B1. Data is shown as mean with SD, n = 3. **B:** Coinfection with $10^6$ cells/ml *E. faecalis* B1 and different cell numbers of *C. albicans*. Data is shown as mean with SD, n = 3. **C:** Coinfection with heat killed *C. albicans* and different *E. faecalis* strains. Data is shown as mean with SD, n = 3. **D:** Coinfection with *C. albicans* and living or heat killed *E. faecalis* strains. Data is shown as mean with SD, n = 4. **A-D**: Coinfection damage and the sum of monoinfection damage was compared for each condition/strain by two-tailored unpaired Student's t-test; p-values < 0.05 are indicated in the graphs; ns: not significant (p > 0.05). **E:** Kinetics of host cell damage during the course of infection. Data is shown as mean with SD, n = 3. Coinfection damage and the sum of monoinfection damage was compared for each time point by two-tailored unpaired Student's t-test corrected for multiple comparisons using the Holm-Sidak method; p-values < 0.05 are indicated in the graphs; ns: not significant (p > 0.05). Damage caused by either *C. albicans* or *E. faecalis* monoinfection was compared across time points by 1-way ANOVA with Tukey's multiple comparison test; only significant (p < 0.05) differences between subsequent time points are indicated with the p-value in the graph. **F:** Impact of preincubation times on coinfection damage (n = 1). **G:** Host cell damage by different haemolytic colonies obtained after conjugation of *E. faecalis* B1 (donor) with *E. faecalis* OG1RF (recipient) (n = 1). **H:** Host cell damage by different non-haemolytic colonies obtained after conjugation of *E. faecalis* B1 (donor) with *E. faecalis* OG1RF (recipient) (n = 1).

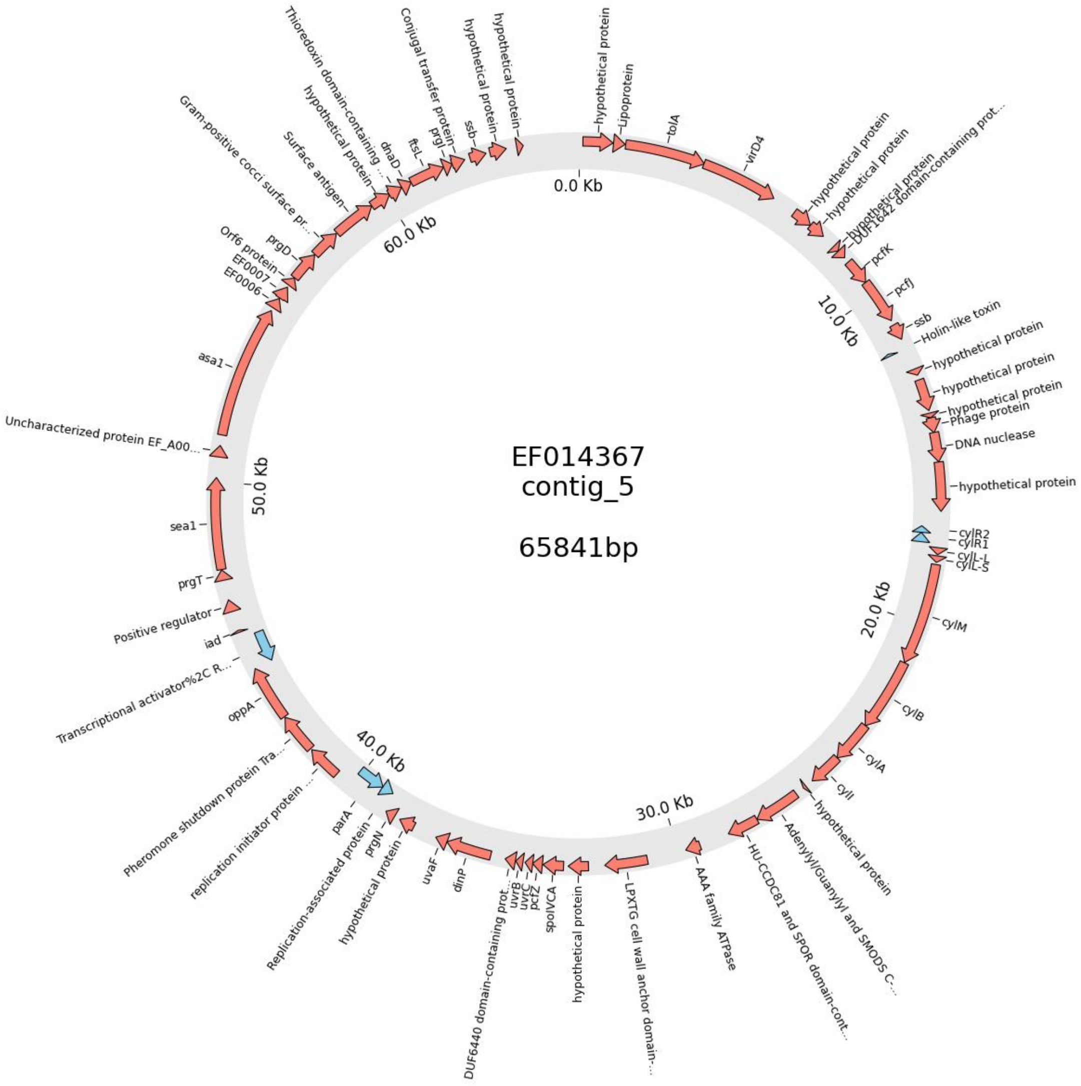

**Fig. S3: Putative conjugative plasmid containing the *cyl* operon in *E. faecalis* B1**

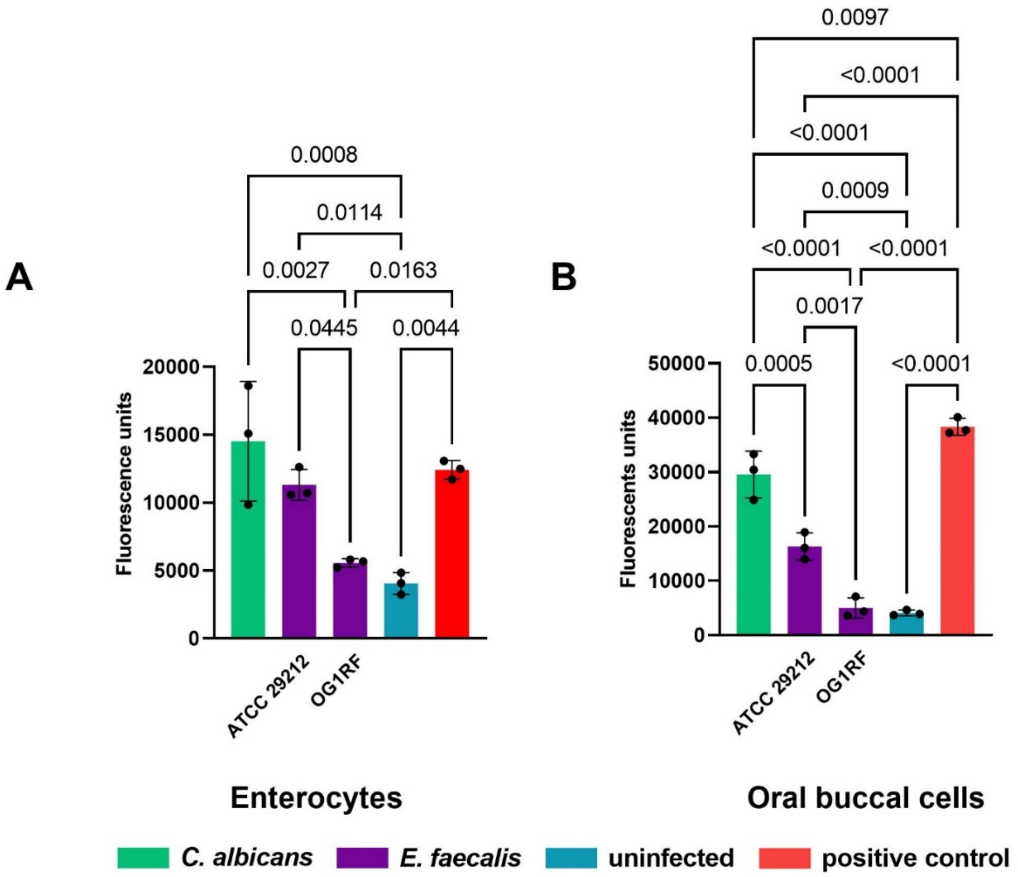

**Fig. S4: Apoptosis of infected enterocytes and oral cells *in vitro***
Enterocytes were infected with $10^6$ cells/ml *C. albicans* SC5314 or $10^5$ cells/ml *E. faecalis* (strain as indicated) at 1 % $O_2$. DMSO was used as positive control. Apoptosis was measured using FAM-DEVD-OPH *in vitro* Apoptosis Detection Reagent and is displayed as Flourescence units. Data from three independent biological replicates is shown as mean with SD, and was analysed by 1-way ANOVA and Tukey's multiple comparisons test. Statistical significance ($p > 0.05$) is indicated by p-values in the graph **A:** Enterocytes were infected for 30 h with *C. albicans* and 24 h with *E. faecalis* before analysis. **B:** Human oral buccal cells (TR146) were infected for 12 h with *C. albicans* and 6 h with *E. faecalis* before analysis.

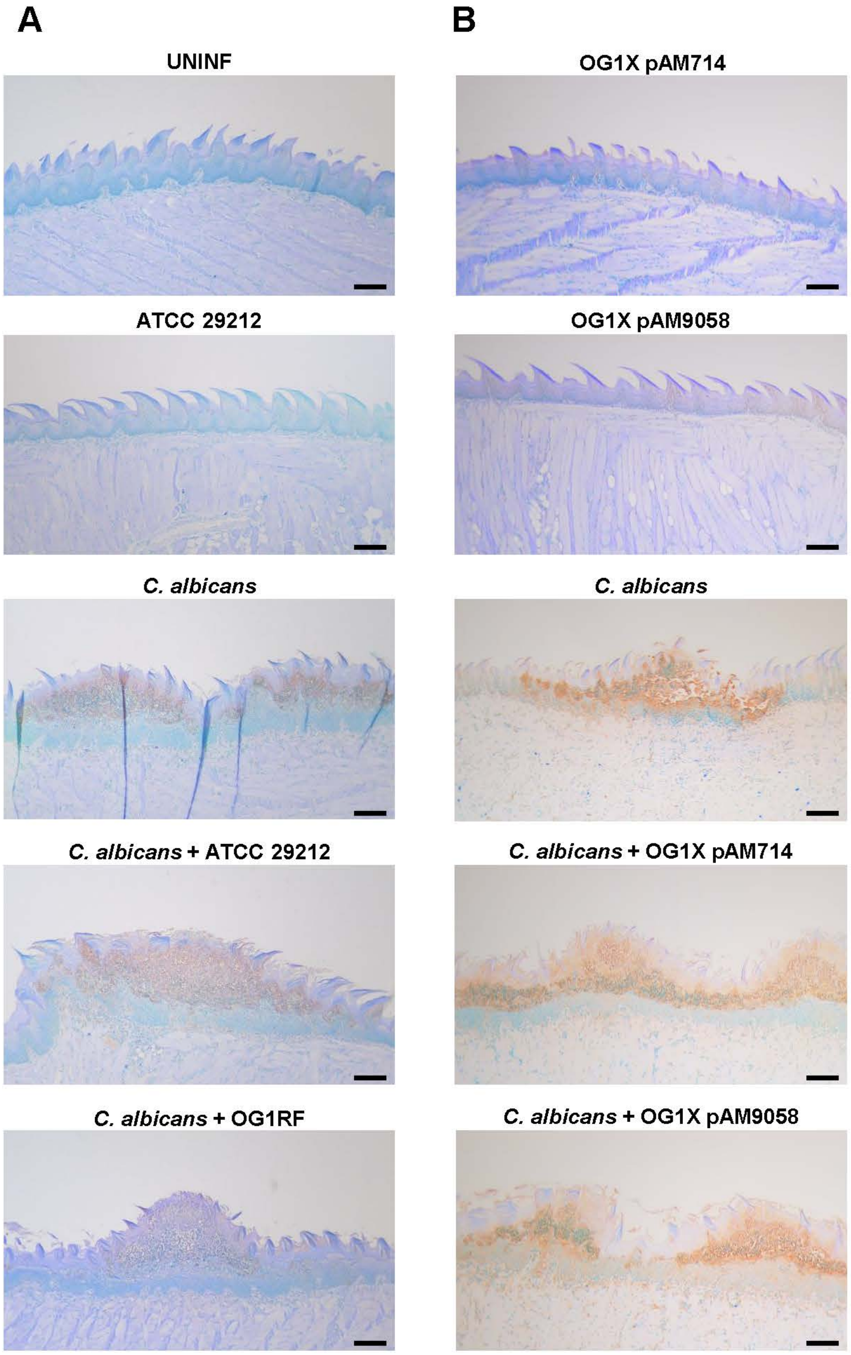

**Fig. S5: Apoptosis in murine tongues during OPC**

Representative micrographs of murine tongue sections stained for apoptosis. Animals infected with *E. faecalis*, *C. albicans*, or both were sacrificed on day 2 after infection. For each animal, three non-consecutive formalin-fixed paraffin-embedded tongue tissue sections (sections from different parts of the tongue) were randomly selected and stained using the ApopTag *in situ* apoptosis detection kit. **A:** Infection with *C. albicans* and/or *E. faecalis* strain ATCC 29212 and OG1RF, respectively. n=5 mice per group. **B:** Infection with *C. albicans* and/or *E. faecalis* strain OG1X pAM714 and OG1X pAM9058, respectively. n=6 mice/group.

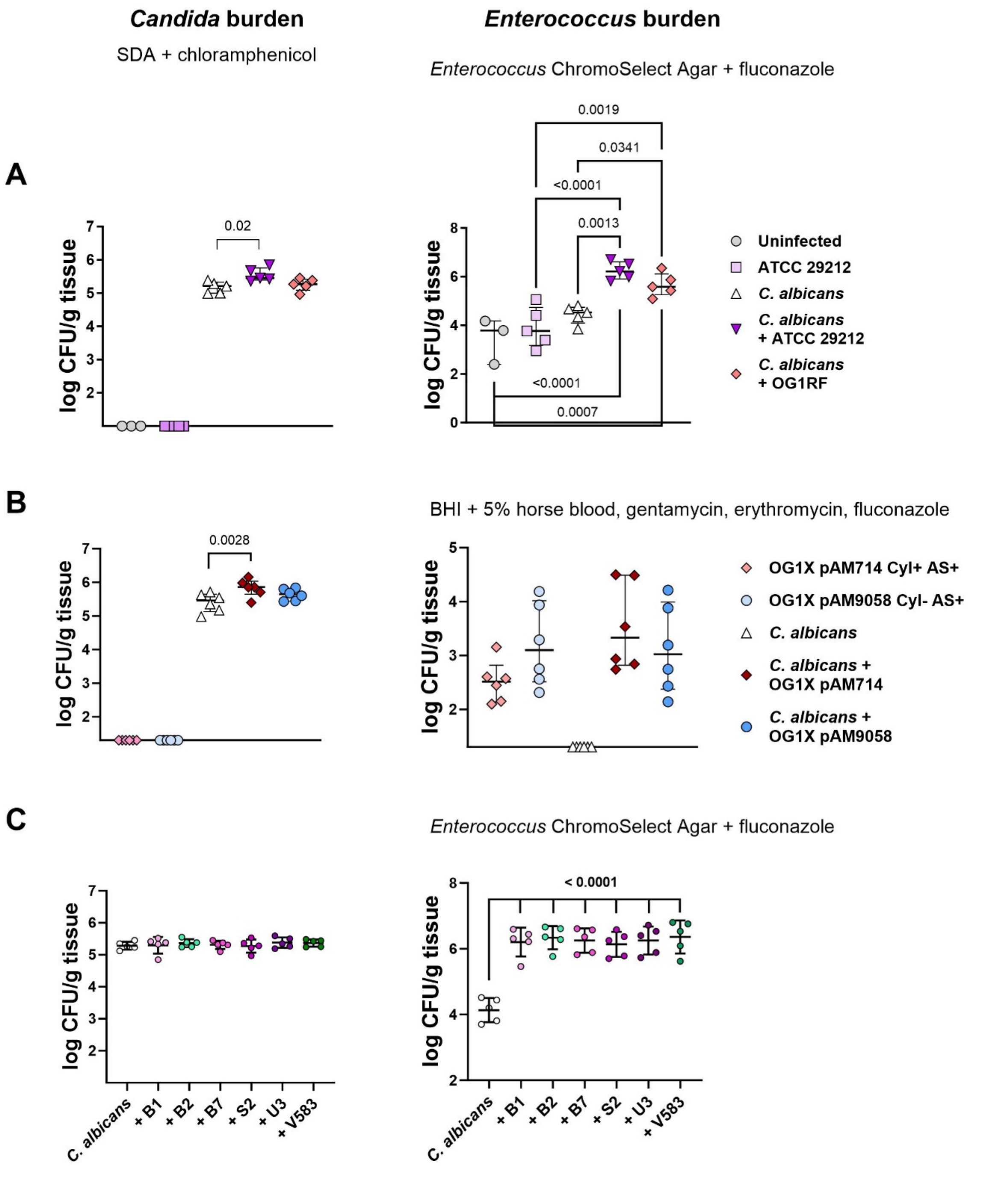

**Figure S6: Microbial burden in murine tongue tissue**
Six to seven week old male C57BL/6J mice were given a subcutaneous injection of 150 mg/kg cortisone acetate at day -1 and +1. C. *C. albicans* infection was induced on day 0 by sublingual application of cotton swaps with *C. albicans* SC5314 for 75 min. Six hours later *E. faecalis* was added to the drinking water for 24 h (2x$10^7$ CFU in 50 ml drinking water). Animals were sacrificed on day 2 after infection. For CFU enumeration organs were homogenized and serial dilutions were plated on selective media as indicated in the above the graphs. Left: *C. albicans*; right: *E. faecalis*; the strains used are indicated in the graphs. Data is presented as individual data points, mean and SD. **A:** Uninfected: Neither *C. albicans* nor enterococci were added. Statistical analysis was performed by one-way ANOVA followed by Tukey's multiple comparison test; for *C. albicans* CFU Uninfected and ATCC 29212 (no *C. albicans* infection) were excluded from the analysis. p values < 0.05 are indicated in the graphs. **B:** Statistical analysis was performed by one-way ANOVA followed by Tukey's

multiple comparison test; for *C. albicans* CFU pAM714 and pAM9058 without *C. albicans* infection were excluded from the analysis. For *E. faecalis* CFU, *C. albicans* (no enterococci added) was excluded from the analysis. p values < 0.05 are indicated in the graphs. **C:** Statistical analysis was performed by one-way ANOVA followed by Tukey's multiple comparison test; p values < 0.05 are indicated in the graphs.

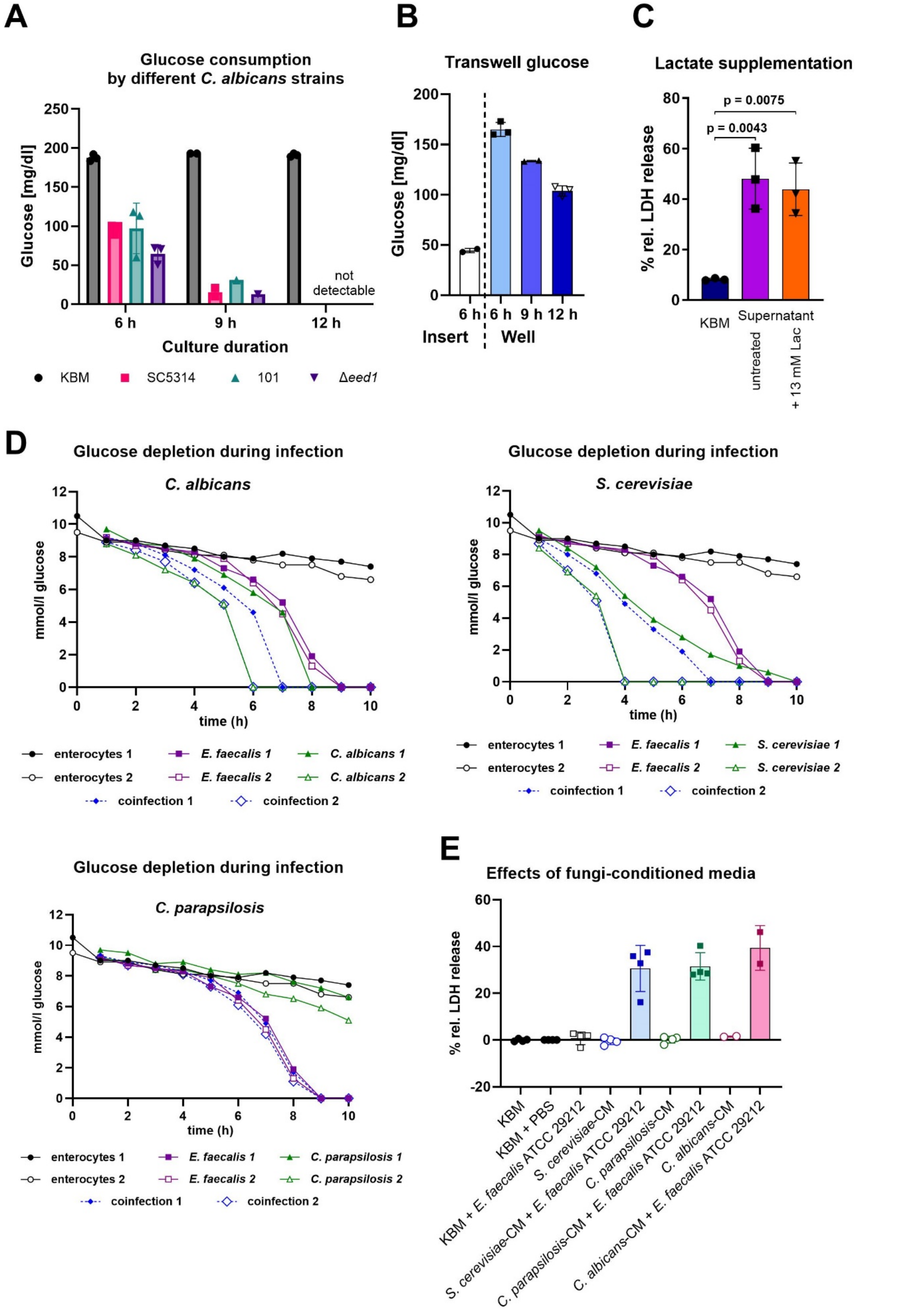

**Fig. S7: Glucose consumption and effects of lactose depletion on enterocyte damage**

**A:** Glucose consumption by different *C. albicans* strains grown in KBM. KBM was inoculated with $10^6$ fungal cells/ml and glucose was quantified using a blood sugar meter (Ascensia Diabetes Care, Germany). **B:** Transwell experiment with 0.4 µm pores size for spatial separation. *C. albicans* SC5314 ($10^6$/ml) was added to the insert at 0 h; *E. faecalis* ATCC 29212 ($10^5$ cells/ml) was added to the well 6 h later. Glucose concentrations were measured as described for A at the indicated time points in the different compartments. **C:** Effect of lactate supplementation (13 mM L-(+)-lactate; 13 mM lac) on enterocyte damage. Enterocytes were infected with $10^5$ cells/ml *E. faecalis* ATCC 29212 (30 h, 1% $O_2$, 5% $CO_2$) using *C. albicans* SC5314 conditioned media produced without enterocytes. Damage was measured by release of host cell lactate dehydrogenase (LDH) into the supernatant, normalised to non-infected controls, and is displayed as percent damage relative to Triton X-100-lysis. Data is shown as mean and SD, n = 3. Statistical analysis was performed by 1-way ANOVA followed by Tukey's multiple comparisons test to compare all groups with each other. Significant differences are indicated by p-values in the graphs. **D:** Glucose concentration in enterocyte supernatants was measured each hour. Host cells were infected with $10^6$ cells/ml fungi, $10^5$ cells/ml *E. faecalis*, or both. Every hour one droplet of supernatant was taken and the glucose measured as described for A. Displayed are two independent measurements for each uninfected enterocytes, bacteria, fungi, and coinfection. **E:** Conditioned media (CM) were prepared by incubating fungi for 30 h (1% $O_2$, 5% $CO_2$) in KBM (without enterocytes). The medium was sterile filtered and used for infection of enterocytes with *E. faecalis* ATCC 29212. Data (n=4, except *C. albicans*: n=2) were analysed by 1-way ANOVA with Dunnetts multiple comparisons test to compare *S. cerevisiae* + *E. faecalis* and *C. parapsilosis* CM + *E. faecalis* with *C. albicans* + *E. faecalis*. The differences were not statistically significant ($p>0.05$).

### Cytolysin expression during coculture

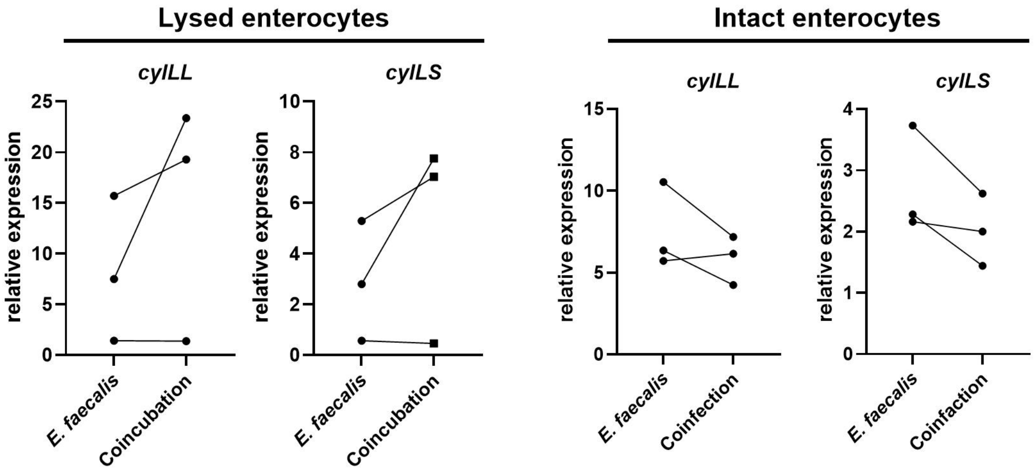

**Fig. S8: Bacterial gene expression during coinfection with different fungi**

Expression of selected *E. faecalis* ATCC 29212 genes, normalised to the housekeeping gene *gyrA*. **A:** Expression of the large (*cylLL*) and small (*cylLS*) cytolysin subunits after 18 h of *E. faecalis* growth. Lysed (left) or cryopreserved (right) enterocytes were added to the medium at the start of coincubation in an equal amount to *C. albicans.* Statistical significance was determined with Wilcoxon matched-pairs signed rank test. No significant differences between the groups were observed ($p > 0.05$).

**Table S1: Virulence properties of the *E. faecalis* strains showing synergistic damage *in vitro***
green: shared features/genes

| Factor | Gene name | *E. faecalis* strains - synergistic | | | | | |
|---|---|---|---|---|---|---|---|
| | | **ATCC 29212** | **B1** | **B7** | **S2** | **S3** | **U3** |
| **Aggregation substance** | as_EF0149 | Y | Y | Y | Y | Y | Y |
| | as_EF0485 | Y | Y | Y | Y | Y | Y |
| | as_prgB_asc10 | Y | Y | Y | Y | Y | Y |
| | asa1 | Y | Y | Y | Y | Y | Y |
| **Capsule** | cpsA | Y | Y | Y | Y | Y | Y |
| | cpsB | Y | Y | Y | Y | Y | Y |
| | cpsC | Y | N | Y | N | N | N |
| | cpsD | Y | N | Y | N | N | N |
| | cpsE | Y | N | Y | N | N | N |
| | cpsF | N | N | Y | N | N | N |
| | cpsG | Y | N | Y | N | N | N |
| | cpsH | Y | N | Y | N | N | N |
| | cpsI | Y | N | Y | N | N | N |
| | cpsJ | Y | N | Y | N | N | N |
| | cpsK | Y | N | Y | N | N | N |
| **Capsule operon type** | | CPS 5 | CPS 1 | CPS 2 | CPS 1 | CPS 1 | CPS 1 |
| **Endocarditis- and biofilm-associated pilus** | ebpA | Y | Y | Y | Y | Y | Y |
| | ebpB | Y | Y | Y | Y | Y | Y |
| | ebpC | Y | Y | Y | Y | Y | Y |
| | srtC | Y | Y | Y | Y | Y | Y |
| **Adhesin** | efaA | Y | Y | Y | Y | Y | Y |
| **Adhesin to collagen** | ace | Y | Y | Y | Y | Y | Y |
| **Enterococcal surface protein** | esp | Y | Y | N | Y | N | Y |

**Table S1: Virulence properties of the *E. faecalis* strains showing synergistic damage *in vitro* - continued**
green: shared features/genes

| **Factor** | **Gene name** | ***E. faecalis* strains - synergistic** | | | | | |
|---|---|---|---|---|---|---|---|
| | | **ATCC 29212** | **B1** | **B7** | **S2** | **S3** | **U3** |
| **Quorum sensing** | fsrA | N | Y | Y | N | Y | N |
| | fsrB | N | Y | Y | N | Y | N |
| | fsrC | Y | Y | Y | Y | Y | Y |
| **Gelatinase** | gelE | Y | Y | Y | Y | Y | Y |
| **Serine Protease** | sprE | Y | Y | Y | Y | Y | Y |
| **Biofilm formation** | bopD | Y | Y | Y | Y | Y | Y |
| **Hyaluronidase** | hyal_EF0818 | N | Y | Y | N | Y | N |
| | hyal_EF3023 | Y | Y | Y | Y | Y | Y |

**Table S2: Virulence properties of the *E. faecalis* strains not showing synergistic damage *in vitro***

green: shared features/genes in synergistic strains

| **Factor** | **Gene name** | ***E. faecalis* strains – non-synergistic** | | | | | | | | | | | | |
|---|---|---|---|---|---|---|---|---|---|---|---|---|---|---|
| | | **ATCC 19433** | **OG1RF** | **V583** | **B2** | **B3** | **B4** | **B6** | **B8** | **B9** | **S1** | **S4** | **U1** | **U2** |
| **Aggregation substance** | as_EF0149 | N | N | Y | Y | Y | N | N | Y | Y | Y | Y | Y | N |
| | as_EF0485 | N | N | Y | Y | Y | N | N | Y | Y | Y | Y | Y | N |
| | as_prgB_asc 10 | N | N | Y | Y | Y | N | N | Y | Y | Y | Y | Y | N |
| | asa1 | N | N | Y | Y | Y | N | N | Y | Y | Y | Y | Y | N |
| **Capsule** | cpsA | N | Y | Y | Y | Y | Y | N | Y | Y | Y | Y | Y | Y |
| | cpsB | N | Y | Y | Y | Y | Y | N | Y | Y | Y | Y | Y | Y |
| | cpsC | N | N | Y | Y | Y | N | N | Y | Y | Y | N | N | N |
| | cpsD | N | N | Y | Y | Y | N | N | Y | Y | Y | N | N | N |
| | cpsE | N | N | Y | Y | Y | N | N | Y | Y | Y | N | N | N |
| | cpsF | N | N | Y | Y | Y | N | N | Y | Y | N | N | N | N |
| | cpsG | N | N | Y | Y | Y | N | N | Y | Y | Y | N | N | N |
| | cpsH | N | N | Y | Y | Y | N | N | Y | Y | Y | N | N | N |
| | cpsI | N | N | Y | Y | Y | N | N | Y | Y | Y | N | N | N |
| | cpsJ | N | N | Y | Y | Y | N | N | Y | Y | Y | N | N | N |
| | cpsK | N | N | Y | Y | Y | N | N | Y | Y | Y | N | N | N |
| **Capsule operon type** | | | CPS 1 | CPS 2 | CPS 2 | CPS 2 | CPS 1 | | CPS 2 | CPS 2 | CPS 5 | CPS 1 | CPS 1 | CPS 1 |

**Table S2: Virulence properties of the *E. faecalis* strains not showing synergistic damage *in vitro* - continued**

green: shared features/genes in synergistic strains

| Factor | Gene name | B: *E. faecalis* strains – non-synergistic | | | | | | | | | | | | |
|---|---|---|---|---|---|---|---|---|---|---|---|---|---|---|
| | | **ATCC 19433** | **OG1RF** | **V583** | **B2** | **B3** | **B4** | **B6** | **B8** | **B9** | **S1** | **S4** | **U1** | **U2** |
| **Endocarditis- and biofilm-associated pilus** | ebpA | N | Y | Y | Y | Y | Y | N | Y | Y | Y | Y | Y | Y |
| | ebpB | N | Y | Y | Y | Y | Y | N | Y | Y | Y | Y | Y | Y |
| | ebpC | N | Y | Y | Y | Y | Y | N | Y | Y | Y | Y | Y | Y |
| | srtC | N | Y | Y | Y | Y | Y | N | Y | Y | Y | Y | Y | Y |
| **Adhesin** | efaA | N | Y | Y | Y | Y | Y | N | Y | Y | Y | Y | Y | Y |
| **Adhesin to collagen** | ace | N | Y | Y | Y | Y | Y | N | Y | Y | Y | Y | Y | Y |
| **Enterococcal surface protein** | esp | N | N | N | N | Y | N | N | Y | Y | Y | Y | N | N |
| **Quorum sensing** | fsrA | N | Y | Y | Y | Y | Y | N | N | Y | N | N | N | Y |
| | fsrB | N | Y | Y | Y | Y | Y | N | N | Y | N | N | N | Y |
| | fsrC | N | Y | Y | Y | Y | Y | N | Y | Y | Y | Y | N | Y |
| **Gelatinase** | gelE | N | Y | Y | Y | Y | Y | N | Y | Y | Y | Y | N | Y |
| **Serine Protease** | sprE | N | Y | Y | Y | Y | Y | N | Y | Y | Y | Y | N | Y |
| **Biofilm formation** | bopD | N | Y | Y | Y | Y | Y | N | Y | Y | Y | Y | Y | Y |
| **Hyaluronidase** | hyal_EF0818 | N | Y | Y | N | Y | Y | N | Y | Y | N | N | Y | Y |
| | hyal_EF3023 | N | Y | Y | Y | Y | N | N | Y | Y | Y | Y | Y | Y |

**Table S3. Cytolysin operon gene distribution of *E. faecalis* strains.**

Strains were streaked from glycerol stocks onto Columbia CNA agar plates containing 5 % horse blood (HBA) and incubated at 21 % or 1 % $O_2$ for 24 h. Strains displaying synergistic coinfection damage are highlighted in pink. The presence of cytolysin genes was evaluated by sequencing genomic DNA or by PCR, complete operons are highlighted in fuchsia. Haemolysis is indicated in orange and ranked from weak, medium, to strong depending on the zone of haemolysis.

| Locus tag gene | DR75_2951 *CylR2* | DR75_2952 *CylR1* | DR75_2953 *CylLL* | DR75_2954 *CylLS* | DR75_2955 *CylM* | DR75_2956 *CylB* | DR75_2957 *CylA* | DR75_2958 *CylI* | β-haemolysis on HBA | | Synergistic *in vitro* |
|---|---|---|---|---|---|---|---|---|---|---|---|
| | | | | | | | | | 21 % O2 | 1 % O2 | |
| ATCC 19433 | N | N | N | N | N | N | N | N | medium | N | N |
| ATCC 29212 | Y | Y | Y | Y | Y | Y | Y | Y | strong | strong | Y |
| OG1RF | N | N | N | N | N | N | N | N | N | N | N |
| V583 | N | N | N | N | N | N | N | N | N | N | N |
| B1 | Y | Y | Y | Y | Y | Y | Y | Y | medium | strong | Y |
| B2 | N | N | N | N | N | N | Y | Y | N | N | N |
| B3 | Y | Y | Y | Y | Y | Y | Y | Y | N | weak | N |
| B4 | N | N | N | N | N | N | N | N | N | N | N |
| B5 | N | N | N | N | N | N | N | N | N | N | N |
| B6 | N | N | N | N | N | N | N | N | N | N | N |
| B7 | Y | Y | Y | Y | Y | Y | Y | Y | weak | medium | Y |
| B8 | Y | Y | Y | Y | Y | Y | Y | Y | N | N | N |
| B9 | Y | Y | Y | Y | Y | N | N | N | N | N | N |
| S1 | N | N | N | N | N | N | N | N | N | N | N |
| S2 | Y | Y | Y | Y | Y | Y | Y | Y | medium | strong | At 10 % $O_2$ |
| S3 | Y | Y | Y | Y | Y | Y | Y | Y | strong | strong | Y |
| S4 | N | N | N | N | N | N | N | N | N | N | N |
| U1 | N | N | N | N | N | N | N | N | N | N | N |
| U2 | N | N | N | N | N | N | N | N | N | N | N |
| U3 | Y | Y | Y | Y | Y | Y | Y | Y | medium | strong | Y |

| Complete cytolysin operon | | | Haemolytic/synergistic phenotype | | |
|---|---|---|---|---|---|
| Y = Yes | N = No | Y = incomplete operon | Y= Yes | Y= Yes | N = No |

**Table S4. Fungal strains used in this work.**

| Strain | Species | Genotype | Reference |
|---|---|---|---|
| SC5314 | *Candida albicans* | prototrophic clinical isolate | (*1*) |
| *eed1* | | *eed1*::FRT/*eed1*::FRT | (*2*) |
| *efg1/cph1* | | *cph1Δ::hisG/cph1Δ::hisG/efg1Δ::hisG/efg1Δ::hisG-URA3-hisG* | (*3*) |
| BWP17 | | *ura3::limm434/ura3::limm434arg4::hisG/arg4::hisGhis1::hisG/his1::hisG* | (*4*) |
| *ece1* | | *ece1::ARG4/ece1::HIS1 RPS1/rps1::URA3* | (*5*) |
| 101 | | prototrophic clinical isolate | (*6*) |
| ATCC 9763 | *Saccharomyces cerevisiae* | | |
| ATCC 2001 | *C. glabrata* | | |
| DSM 4237 | *C. parapsilosis* | | |

**Table S5.** ***Enterococcus faecalis*** **strains used in this work.**

| Strain designation | Phenotype/Origin | Resistance markers* | CFUs/ml at $OD_{600}$ = 1 exponential growth | Reference |
|---|---|---|---|---|
| ATCC 19433 | type strain | | $7.79x10^7$ | |
| ATCC 29212 | vancomycin-sensitive, representative control strain for clinical and laboratory experiments | | $2.01x10^8$ | |
| V583 | first vancomycin-resistant clinical isolate reported in the USA; bloodstream isolate | *van* | $1.60x10^8$ | (*7*) |
| OG1RF | derivative of OG1 selected for spontaneous resistance to rif and fus; plasmid free | *rif, fus* | $3.25x10^8$ | (*8*) |
| OG1RF Cyl+ | OG1RF conjugated with B1, haemolytic | *rif, fus* | $3.25x10^8$ | this work |
| OG1RF Cyl- | non-haemolytic colony obtained after conjugation of OG1RF with B1 | *rif, fus* | $3.25x10^8$ | this work |
| OG1X | gelatinase-negative derivative of OG1-10 (OG1-derivative, streptomycin-resistant); plasmid free | *str* | $5.18x10^8$ | (*9*) |
| pAM714 | OG1X derivative carrying pAM1::Tn*917* (pAM714) $Agg^+$ $Cyl^+$; | *str, erm* | $5.82x10^8$ | (*10*) |
| pAM944 | OG1X derivative carrying pAM1::Tn*917* (pAM944) $Agg^-$ $Cyl^+$ | *str, erm* | $6.38x10^8$ | (*11*) |
| pAM9058 | OG1X derivative carrying pAM1::Tn*917* (pAM9058) $Agg^+$ $Cyl^-$ | *str, erm* | $8.63x10^8$ | (*12*) |
| B1 | clinical blood isolates | | $6.24x10^8$ | UKJ |
| B2 | | | $3.68x10^8$ | |
| B3 | | | $5.30x10^8$ | |
| B4 | | | $5.62x10^8$ | |
| B5 | | | $5.30x10^8$ | |
| B6 | | | $6.89x10^8$ | |
| B7 | | | $3.87x10^8$ | |
| B8 | | | $2.44x10^8$ | |
| B9 | | | $2.69x10^8$ | |
| S1 | clinical stool isolates | | $5.62x10^8$ | |
| S2 | | | $7.28x10^8$ | |
| S3 | | | $3.04x10^8$ | |
| S4 | | | $7.12x10^8$ | |
| U1 | clinical urine isolates | | $4.39x10^8$ | |
| U2 | | | $3.01x10^8$ | |
| U3 | | | $4.21x10^8$ | |

| | | | |
|---|---|---|---|
| B1 cure (Cyl-) | B1 derivative from plasmid curing; non-hemolytic | $6.24x10^8$ | this work |
| B1 cure (Cyl+) | B1 derivative from plasmid curing; hemolytic | $6.24x10^8$ | this work |

* Resistance markers: *van*: vancomycin; *rif*: rifampicin; *fus*: fusidic acid; *str*: streptomycin; *erm*: erythromycin
